# Accurate and Efficient Estimation of Local Heritability using Summary Statistics and LD Matrix

**DOI:** 10.1101/2023.02.08.527759

**Authors:** Hui Li, Rahul Mazumder, Xihong Lin

## Abstract

Existing SNP-heritability estimation methods that leverage GWAS summary statistics produce estimators that are less efficient than the restricted maximum likelihood (REML) estimator using individual-level data under linear mixed models (LMMs). Increasing the precision of a heritability estimator is particularly important for regional analyses, as local genetic variances tend to be small. We introduce a new estimator for local heritability, “HEELS”, which attains comparable statistical efficiency as REML (*i.e*. relative efficiency greater than 92%) but only requires summary-level statistics – Z-scores from the marginal association tests plus the empirical LD matrix. HEELS significantly improves the statistical efficiency of the existing summary-statistics-based heritability estimators– for instance, HEELS produces heritability estimates that are more than 3-fold and 7-times less variable than GRE and LDSC, respectively. Moreover, we introduce a unified framework to evaluate and compare the performance of different LD approximation strategies. We propose representing the empirical LD as the sum of a low-rank matrix and a banded matrix. This approximation not only reduces the storage and memory cost of using the LD matrix, but also improves the computational efficiency of the HEELS estimation. We demonstrate the statistical efficiency of HEELS and the advantages of our proposed LD approximation strategies both in simulations and through empirical analyses of the UK Biobank data.

## Introduction

In the last decade, advances in biotechnology have enabled estimation of genetic variance contributed by genotyped variants without requiring assumptions about the shared environmental effects^1–6^. These methods estimate the so-called SNP-heritability 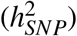, defined as the proportion of phenotypic variance due to causal variants tagged by genotyped variants, and have led to critical insights into the genetic architectures of complex traits and diseases^7^. The existing 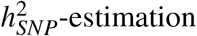 methods can be broadly categorized as either based on individual-level genotype and phenotype data^1,2,8–11^ or based on summary statistics from genome-wide association studies (GWAS)^3,4,6,12^.

Estimators that use individual-level data are generally more precise, but the applicability of these methods can be limiting due to privacy concerns and data-sharing restrictions. Heritability estimation methods that are based on GWAS summary statistics have gained popularity in recent years, but an important drawback of these approaches is low statistical efficiency^11,13^. For instance, studies have shown that the empirical variance of the estimates from LD-score regression (LDSC) is much larger than that of a REML-based estimator^14,15^. Other methods, such as Generalized Random Effects (GRE) and Randomized HE-regression (RHE-reg), are also less precise than REML based on simulations^10,11^, To address these limitations, we introduce <u>H</u>eritability <u>E</u>stimation with high <u>E</u>fficiency using <u>L</u>D and association <u>S</u>ummary Statistics (HEELS) – an accurate and statistically efficient estimator of local 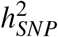 that only requires summary-level statistics.

Our work has two main contributions to the literature of 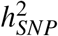 estimation. First, we propose an iterative procedure that uses summary statistics to solve the REML-score equations by transforming the Henderson’s algorithm for estimating the variance components in linear mixed models^16,17^. The summary statistics required by HEELS includes the marginal association statistics from GWAS and the correlation statistics estimated in-sample (*i.e*. empirical covariance matrix of the genotypes, “LD”). We show both analytically and through extensive simulations that the summary-statistics-based HEELS estimator attains comparable level of statistical efficiency or precision as the conventional individual-level-data-based REML estimators, such as GREML^1^ and BOLT-REML^2^. The relative efficiency of HEELS is significantly higher than that of the state-of-the-art summary-statistics-based methods, i.e., the estimates from HEELS have much smaller variance. Our method is applicable and most suitable to local heritability estimation, as the genetic variance at a specific region tends to be small and thus its statistical efficiency is particularly important. Second, we introduce a novel low-dimensional representation of the LD as the sum of a banded and a low-rank matrix (“Banded + LR”). This structure helps provide a unified framework where many of the existing LD approximations can be viewed as special cases. We use an optimization approach to solve for the best representation of LD with this “Banded + LR” structure, and evaluate the performance of different low-dimensional approximations in the context of 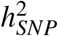 estimation. Compared to existing methods, our proposed decomposing approach not only achieves greater approximation accuracy of the LD matrix, measured in the matrix spectral norm, but also yields heritability estimates with smaller bias.

We applied HEELS to estimate the local SNP-heritability of 30 complex traits and diseases in the UK Biobank (UKB). Consistent with our findings in the simulations, the estimates from HEELS are highly concordant with the estimates from REML. HEELS also attains the highest statistical efficiency compared to the other summary-statistics-based methods. We used the local 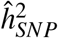 estimates from HEELS to contrast the genetic architectures of traits, and found different distribution patterns of local genetic variance, consistent with previous studies^18,19^. We also used HEELS to prioritize genetic regions that are enriched for related traits (*e.g*. lipid traits, blood-cell and leukocyte traits). The loci we identified not only corroborate known mechanisms of disease-association, but also yield novel insights into the shared genetic basis among correlated traits, pointing to putative pleiotropic hotspots.

Several recently developed methods are related to our work. Here we highlight the their similarities and distinctions compared to HEELS. High-Definition Likelihood (“HDL”) has a similar aim as our work in terms of improving statistical efficiency^20^ (although the method was proposed for genetic correlation estimation, it necessarily computes heritability as an intermediate step). Notably, our estimator was derived based on the marginal likelihood of the phenotypes (although the estimation does not require accessing individual-level data) while HDL uses the likelihood of the Z-statistics. In terms of LD approximation, our proposed banded + low-rank representation of the LD differs from the truncated SVD approach used by HDL. In simulations, we observed bias in HDL’s estimates of heritability, which has been reported in the benchmarking results of another study^21^. Therefore, we do not include HDL in our comparisons among *unbiased* estimators. Another recently published work introduces method called “LDAK-GBAT”, using the same likelihood functions as ours^22^. However, LDAK-GBAT focuses on gene-based association testing rather than heritability estimation. The paper does not report or analyze the statistical efficiency of the heritability estimator. LDAK-GBAT also uses a permutation procedure which inherently requires individual-level data, whereas HEELS solely relies on summary-level statistics for both heritability estimation and standard error calculation. Both HDL and LDAK-GBAT maximize their respective likelihood functions directly using a Newton-Raphson type algorithm, whereas our iterative algorithm operates on the score equations and involves *closed-form* updating equations. Finally, we note that a parallel work – SuSiE-RSS – also exploits the sufficient statistics of the likelihood function to enable “dual” algorithms that are applicable to both individual-level data and summary-level statistics^23^. In other words, the relationship between SuSiE and SuSiE-RSS is akin to the relationship between GREML and HEELS. Nevertheless, SuSiE-RSS is designed for fine-mapping and it performs variable selection under the Bayesian framework, which is distinct from HEELS.

The literature of SNP-heritability estimation is quite rich. From a computational perspective, advances in scalable heritability estimation algorithms have enabled existing estimators, such as those from REML and HE-regression, to be applied to biobank-sized datasets^2,11,24^. From a modeling perspective, researchers have made different assumptions about the distribution of effect sizes^9,25,26^. Although efforts have been made to reconcile these modeling inconsistencies^13,25,27^, debates remain unresolved as to which assumptions are more fitting and realistic. We emphasize that the particular focus of our study is to improve the *statistical efficiency* of heritability estimators that are based on summary-level statistics. It is out of the scope of our current work to compare across all or most of the existing heritability estimation methods, nor to evaluate the suitability of any particular heritability model. We choose LMM as the simplest and most widely understood model to demonstrate the merits of our approach both conceptually and empirically. We hope our work can encourage development of methods that produce more precise heritability estimates under potentially different modeling frameworks.

## Results

### Overview of HEELS

HEELS uses summary-level statistics, including marginal association statistics and the LD matrix, to estimate heritability by iteratively solving the REML score equation^17^. HEELS provides heritability estimates that are as precise as those based on REML-based estimates that use individual-level data, such as GREML^1^. We first introduce some notations and define 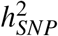 using the linear mixed model based on individual-level data.

Let **y** be a length-*n* vector that denotes the phenotypes of *n* samples. Denote by **X** ∈ ℝ^*n*×*p*^ the genotype matrix of *n* individuals based on *p* markers or SNPs. We standardize **X** and **y** such that the variance of the phenotype is 1 and the variance of each marker-specific genotype vector is 1/*p*, or *diag*(**X**^⊤^**X**/*n*) = 1/*p*. Let **S** and **R** denote the the marginal association statistics and the in-sample LD matrix, *i.e*. 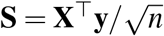 and **R** = **X**^⊤^**X**/*n*. We use an additive genetic model for the phenotypes as **y** = **X***β* + *ε*, where ***β*** is a *p* × 1 vector assumed to follow 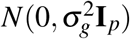, and *ε* is a length-*n* vector distributed as 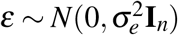. Under these assumptions, **y** ~ *N*(0, **V**), where the variance-covariance matrix is 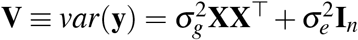. Under this model, SNP-heritability is defined as 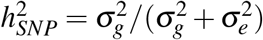 (**Online Methods**).

Suppose we do not have access to individual-level data **X** and **y**, and are only provided with the marginal association statistics **S** = **X**^⊤^**y** and the LD matrix **R** = **X**^⊤^**X**. We show that the REML score equations for 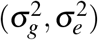 can be solved using these summary-level statistics **S** and **R** only, by applying the Sherman-Woodbury matrix identity to the Anderson’s algorithm for solving the variance components (**Supplementary Notes**). HEELS iterates between updating the Best Linear Unbiased Predictor (BLUP) estimates^17^ of the joint effect sizes 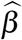 and updating the variance component estimates 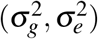 until convergence. Let the superscript^(*t*)^ denote the value of the parameter at iteration *t*. The HEELS estimation procedure is as follows.

1. Update the BLUP joint effect size estimates using:

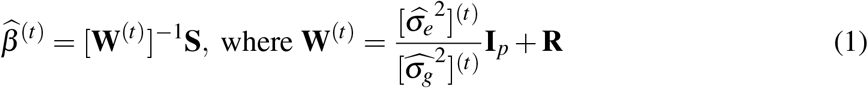
2. Update 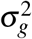 using:

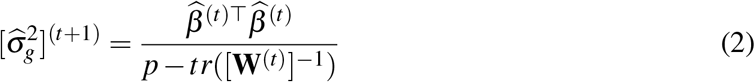
3. Update 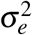 using:

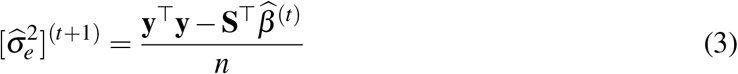

We initialize 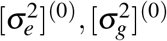 with some random values between 0 and 1, and repeat step 1-3 below until convergence or when the change in *ℓ_HEELS_* between two consecutive iterations is sufficiently small.

We provide derivation details on the updating equations (1)-(3) and further explain the intuition behind these expressions in the **Supplementary Notes**. Notably, the algorithm above does not use the raw individual-level data **X** or **y**, but only utilizes summary statistics **S** and **R**, which are sufficient statistics for the variance component parameters in our model. When sample variance **y**^⊤^**y**/*n* is known, implementing the procedure above is straightforward. When it is not known, we can use the Z-scores, approximate **y**^⊤^**y**/*n* by 1 and rescale 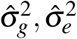 at each iteration accordingly. Similar approximation strategies have been adopted by other summary-statistics-based methods, such as SuSiE-RSS.^23^

For a given model, the maximum-likelihood estimator has the minimal variance among unbiased estimators^28^. The *statistical efficiency* of a heritability estimator is calculated using the ratio of the variance of the REML estimator and the variance of the estimator in comparison^29^. The primary reason for the high statistical efficiency of HEELS is that the estimating equations of 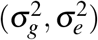 solved by HEELS are identical to the REML score equations of 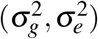^17^. The asymptotic variance of the variance components can be derived using the likelihood theory. Importantly, the variance estimator can also be re-experssed using summary statistics, **S**, **R**, only. We apply the multivariate Delta method to obtain a plug-in estimator of the variance of 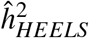 (**Online Methods**).

### Accuracy and statistical efficiency of the HEELS estimator

To evaluate the performance of the HEELS heritability estimator, we performed simulations using the real genotype array data of 332,430 unrelated British white individuals in the UK Biobank. We selected GREML, LDSC, GRE and HESS as the representative 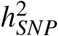 to be compared with HEELS, and do not include other methods in the comparisons due to our focus on *statistical efficiency*. In the **Supplementary Notes**, we provide a more thorough review of the existing 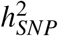 approaches, with their key advantages and limitations summarized in **Supplementary Table** 1, and explain our consideration in selecting the estimators for comparisons.

We first simulated quantitative phenotypes using the LMM assumption (*i.e*. also used by GREML and LDSC), where the causal effect sizes of all markers follow a normal distribution and contribute to the genetic variance equally. We found that HEELS is unbiased in finite samples and attains high statistical efficiency. In particular, we found a high degree of concordance between the estimates from REML-based estimators (such as GREML and BOLT-REML) which use individual-level data and those from HEELS which only relies on summary statistics (**Figure 1)**. When the full UKB data is used, the relative statistical efficiency of HEELS is as high as 99.44%, whereas GRE and LDSC have much lower RE’s of 19.81% and 8.68%, respectively (**Table 1)**.

**Figure 1.**
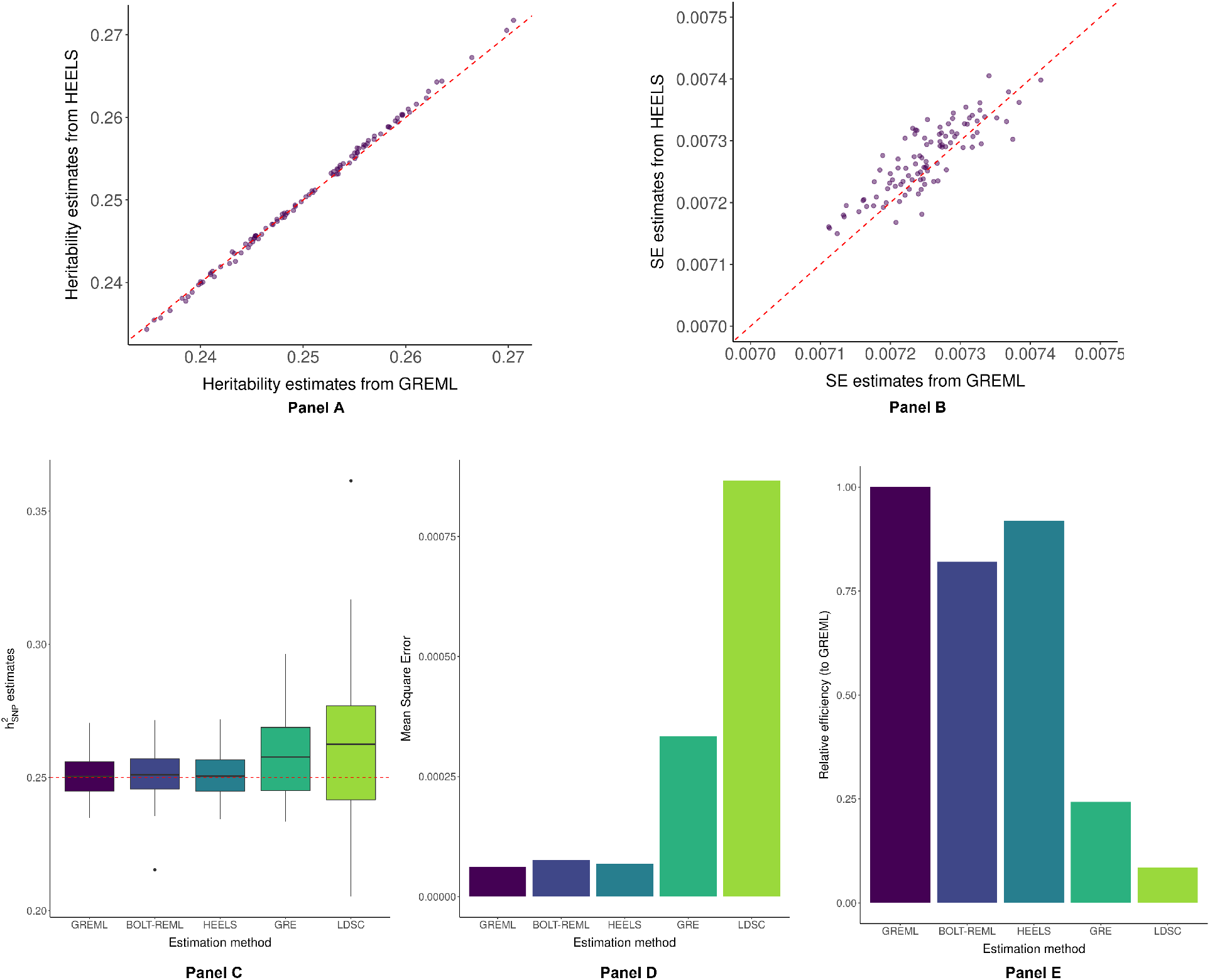
Comparison of the performance of HEELS with different methods using simulation studies. Simulated phenotypes using real genotypic data from the UK Biobank, array SNPs on chromosome 22 with MAF > 0.01 (*n* = 30,000, *p* = 9,205). **Panel A**: Local 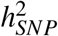 estimates from HEELS using summary statistics vs GREML using individual-level data. Red-dotted line: *y* = *x*. **Panel B**: Analytical SE estimates from HEELS vs GREML. Red-dotted line: *y* = *x*. **Panel C**: Distribution of the 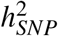 estimates using different methods: GREML and BOLT-REML use individual level data; HEELS, GRE and LDSC use summary statistics. Red-dotted line: true 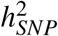 of 0.25. The lower and upper hinges correspond to the first and third quartiles (the 25th and 75th percentiles). The upper (lower) whisker extends from the hinge to the largest (smallest) value no further than 1.5 × *IQR* from the hinge (where IQR is the inter-quartile range). Data beyond the end of the whiskers are called “outlying” points and are plotted individually. **Panel D**: MSEs of the 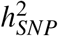 estimates using different methods. **Panel E**: Relative efficiency of different methods compared to GREML.

**Table 1.** Simulation results of the relative statistical efficiencies of different summary-statistics based 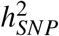 estimates (HEELS, GRE, HESS) compared with individual-level data based GREML. Sparsity and sample sizes are varied across settings. REML estimates are computed using GCTA for small sample (30k), and BOLT-LMM for large samples (40k, 140k, 332k). Statistical relative efficiency: ratio of the variance of REML and the variance of an estimator. Polygenicity: The proportion of the markers that are causal.

| Polygenicity | Sample size | Empirical variance of heritability estimates |  |  |  | Relative efficiency (to REML) |  |  |
| --- | --- | --- | --- | --- | --- | --- | --- | --- |
|  |  | REML | HEELS | GRE | LDSC | HEELS | GRE | LDSC |
| 1 | 332k | 1.43E-05 | 1.44E-05 | 7.24E-05 | 1.65E-04 | 99.44% | 19.81% | 8.68% |
| 1 | 140k | 2.15E-05 | 2.20E-05 | 6.53E-05 | 1.77E-04 | 97.82% | 32.94% | 12.17% |
| 1 | 40k | 4.05E-05 | 4.07E-05 | 1.32E-04 | 5.45E-04 | 99.54% | 30.76% | 7.43% |
| 1 | 30k | 6.19E-05 | 6.74E-05 | 2.55E-04 | 7.32E-04 | 91.91% | 24.25% | 8.46% |
| 0.5 | 30k | 6.80E-05 | 7.26E-05 | 2.58E-04 | 8.71E-04 | 93.69% | 26.38% | 7.81% |
| 0.2 | 30k | 9.35E-05 | 1.00E-04 | 2.98E-04 | 8.89E-04 | 93.16% | 31.42% | 10.52% |
| 0.1 | 30k | 1.16E-04 | 1.29E-04 | 4.39E-04 | 8.91E-04 | 90.17% | 26.41% | 13.02% |

LDSC’s lack of statistical efficiency has been observed in previous studies^12,14,20,30^. We provide a stylized argument to contrast the statistical efficiency of LDSC and HEELS under the framework of moment-matching methods^14^ (**Supplementary Notes**). Briefly, the score equations solved by HEELS coincide with the estimating equations of the most efficient generalized method-of-moment estimator. Since LDSC can be viewed as a method-of-moment estimator with sub-optimal weights, its statistical efficiency is expected to be lower than that of HEELS. In simulations, we found that GRE and HEELS are both unbiased regardless of sample size, but GRE is less statistically efficient that HEELS (**Supplementary Figure** 3). The variability of the GRE estimates is particularly high when sample size is small, as expected from the theory of GRE^10^. Even when the full UKB is used, the variance of 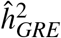 is still five times as large as that of 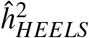. We also compared the performance of HEELS with that of HESS, which is a state-of-the-art method for local heritability estimation^12^. We found that the HESS estimator is sensitive to the degree of LD regularization applied through the truncated SVD (**Supplementary Figure** 5), a phenomenon that has been reported in the original paper. In contrast, HEELS is unbiased when the full LD matrix is used, and we explicitly minimize the bias in the heritability estimate when we approximate the LD (see below for details). We note that the analytic form of the HESS estimator closely resembles that of the GRE estimator, despite their disparate modeling frameworks, as both GRE and HESS can be viewed as principal-component regression (PCR)-based estimators. In simulations, we indeed observed that the HESS estimates are closer to the GRE estimates as the amount of LD regularization decreases (**Supplementary Figure** 5), matching the theory which implies an alignment of 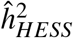 and 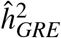 when LD is estimated in-sample without regularization.

Next, we considered relaxing the infinitesimal assumption and altering the sparsity of the causal genetic effects. Although this violates the assumptions used in LMM, we consider it an important setting to evaluate the robustness of our estimator, because it is plausible that only a small fraction of the variants have non-zero effects, as has been assumed by many existing methods^31,32^. In simulations, we found that HEELS remains unbiased and statistically efficient in these mis-specified scenarios (**Table 1)**. Sparsity of the causal effects generally increases the variability of the heritability estimates, but the RE of HEELS remains the highest among summary-statistics-based methods (**Supplementary Figure** 1). Across settings of various levels of polygenicity and sample sizes, HEELS leads to an increase in precision that is equivalent to increasing the GWAS sample size at least 3 times or 7 times, respectively (**Supplementary Figure** 2). Together, our results are consistent with previous works which showed that REML is unbiased even when the modeling assumption is wrong (in terms of point-normal effects)^10,25^, further illustrating the affinity of HEELS and REML. Our findings also corroborate some of the recent theoretical advances in REML estimator analyses, which prove that REML is consistent under non-infinitesimal models^33^ and that sparse architecture will result in a larger asymptotic variance^34^

Finally, we evaluated the Type I error rates controlled by different methods. We found that the standard error of HEELS is well-calibrated regardless of sample size when the model is correctly specified (**Supplementary Figure** 6). The standard errors reported by BOLT-REML using individual-level data generally lead to the correct coverage but can be anti-conservative when sample size is small. The standard errors from GRE can result in under-coverage, as has been reported (see Supplementary Table 4 of Hou *et al*.^10^). When the model is mis-specified (in terms of point-normal effect sizes), we observe under-coverage for all LMM-based estimators. Nevertheless, the calibration of HEELS is still better than GRE and LDSC, and is most comparable to REML across settings (**Supplementary Figure** 7).

### A unified framework to compare LD approximations

It is a well-known challenge in statistical genetics that the LD matrix is expensive to store and compute with. While several low-dimensional representations of the LD matrix have been proposed and used in the literature (*e.g*. banding^35–37^, truncated SVD^12,38^ and low-rank approximation^20,30^), no previous study has systematically compared the performance of different LD approximation strategies. The impact of LD approximations on heritability estimation is also unclear. Our goal is to construct a low-dimensional representation of the LD matrix that can reduce the storage and computational cost of heritability estimation without incurring large loss of accuracy or efficiency in the 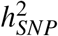 estimates.

To this end, we propose representing the in-sample LD matrix, **R**, as the sum of a banded matrix and a low-rank matrix (“Banded + LR”),

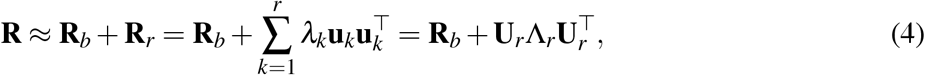

where **R**_*b*_ is a banded matrix with central bandwidth *b*; **R**_*r*_ is a low-rank matrix with rank *r*; *λ_k_* and **u**_*k*_ (also diagonal entries of Λ_*r*_ and columns of **U**_*r*_) are the top-*r* eigenvalues and eigenvectors of **R**_*r*_. We consider a total of six strategies to decompose or represent the empirical LD using this “Banded + LR” structure (**Table 2)**, all of which take the form of the expression in equation (4), but differ in terms of 1) whether the banded component is a diagonal matrix, in which case the approximation becomes a spiked covariance matrix^39^, and 2) whether the banded and low-rank components are constrained to be positive semi-definite (PSD).

**Table 2.**
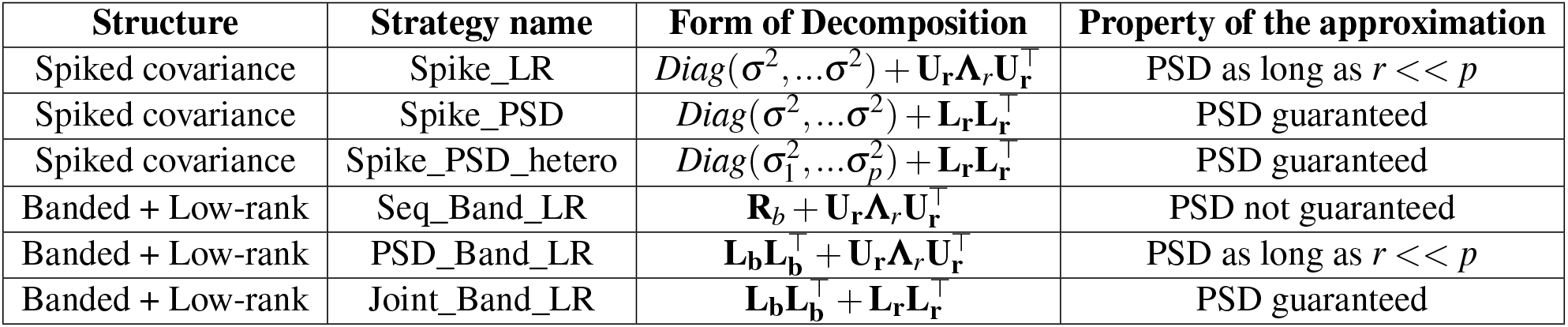
Summary of the proposed LD approximation methods. Six methods are proposed to approximate the LD matrix using either 1) a the spiked covariance matrix (Spike_LR, Spike_PSD, Spike_PSD_hetero) or 2) the sum of a banded and a low-rank matrix (Seq_Band_LR, PSD_Band_LR, and Joint_Band_LR). See **Online Methods** and **Supplemental Notes** for further explanations. PSD: positive semi-definite. *b*: bandwidth of the banded component; *r*: rank of the low-rank component. {**L**, **U**, **Λ**}: Cholesky factor, eigenvector and eigenvalue of the target matrix. Subscript of **L**, **U**, **Λ** represents the component, *b* for banded or *r* for low-rank.

| Structure | Strategy name | Form of Decomposition | Property of the approximation |
| --- | --- | --- | --- |
| Spiked covariance | Spike_LR | $Diag(\sigma^2, \dots, \sigma^2) + \mathbf{U}_r \mathbf{\Lambda}_r \mathbf{U}_r^\top$ | PSD as long as $r \ll p$ |
| Spiked covariance | Spike_PSD | $Diag(\sigma^2, \dots, \sigma^2) + \mathbf{L}_r \mathbf{L}_r^\top$ | PSD guaranteed |
| Spiked covariance | Spike_PSD_hetero | $Diag(\sigma_1^2, \dots, \sigma_p^2) + \mathbf{L}_r \mathbf{L}_r^\top$ | PSD guaranteed |
| Banded + Low-rank | Seq_Band_LR | $\mathbf{R}_b + \mathbf{U}_r \mathbf{\Lambda}_r \mathbf{U}_r^\top$ | PSD not guaranteed |
| Banded + Low-rank | PSD_Band_LR | $\mathbf{L}_b \mathbf{L}_b^\top + \mathbf{U}_r \mathbf{\Lambda}_r \mathbf{U}_r^\top$ | PSD as long as $r \ll p$ |
| Banded + Low-rank | Joint_Band_LR | $\mathbf{L}_b \mathbf{L}_b^\top + \mathbf{L}_r \mathbf{L}_r^\top$ | PSD guaranteed |

Our approximation provides a unified framework for analyzing and evaluating the performance of various LD approximations, as most of the existing LD approximation strategies can be viewed as special cases of decomposition (4). For instance, regularization of the LD matrix through the truncated SVD is equivalent to using “Banded + LR” by setting *b* = 0; the most common approximation of the LD matrix only involves the banded part, so it corresponds to “Banded + LR” by setting *r* = 0. The motivation behind our proposed “Banded + LR” structure (*b* ≠ 0, *r* ≠ 0) is twofold. On one hand, because correlations between two genetic markers are typically induced by physical proximity, the vast majority of the non-zero elements of LD matrix lie on the central band. On the other hand, we want to retain the in-sample correlation structure outside of the central band, because oftentimes there are appreciable non-zero off-central-band elements in finite samples.

We adopt an optimization approach to solve for the best representation of LD with the “Banded + LR” structure, minimizing 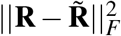, where 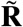 is the working approximating matrix in the form specified in the third column of **Table 2**. We explain the distinctions between different LD approximations in **Online Methods** and provide details on their respective estimation procedures in **Supplementary Table 2**. Briefly, the “Seq_Band_LR” strategy first bands the LD matrix and then performs low-rank decomposition on the residual off-banded matrix; the “PSD_Band_LR” strategy first approximates the banded component of the LD matrix using a PSD matrix and then performs low-rank decomposition on the residual matrix; the “Joint_Band_LR” strategy jointly approximates the banded and the low-rank components using PSD matrices. The PSD assumption can not only reduce the computational cost to solve for the optimal low-dimensional LD representation, but can also improve the computational efficiency of 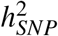 estimation in HEELS (**Supplementary Notes**). We derived the asymptotic variance of our HEELS estimator so that the additional variance incurred by the approximation can be quantified without access to the full LD (**Supplementary Notes**).

### Efficacy of the banded + low-rank representation

In simulations, we found that our proposed “Banded + LR” representations approximate the original full LD with higher accuracy (**Supplementary Figure** 8) and lead to less biased 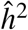 than the existing methods (**Supplementary Figure** 9). We attribute the efficiency of our proposed “Banded + LR” representation to two aforementioned motivating factors, namely, retaining the central band of the covariance matrix while exploiting the signals in the residual off-central-band part of the LD matrix.

Relative to the HEELS estimator that employs the full LD (“Exact_LD”), the runtime of HEELS estimation using the “Seq_Band_LR”, “PSD_Band_LR” or “Joint_Band_LR” approximation is on average 9.52%, 42.61% and 47.37% less. The improvement of computational efficiency pertains to our application of the Woodbury formula to the low-rank matrices, which helps circumvent the need of direct matrix inversions in HEELS estimation. For certain fine-tuned approximation settings (*e.g*. “Seq_Band_LR” with *b* = 600 and *r* = 650), the bias in heritability estimate based on cross-validation is even smaller than that when the full LD is used, and MSE is 3.95% lower even though variance is slightly higher. Overall, across the hyperparameter settings, our proposed “Banded + LR” representation improves the computational efficiency of HEELS, without incurring much loss of accuracy and efficiency (**Table 3)**.

**Table 3.** Comparison of the performance of different LD approximation methods from hyperparmaeter tuning. Top table represents the average performance across hyperparameter settings, with *b* varying from 300 to 600 in increments of 100, and *r* varying from 300 to 800 in increments of 50. The second column shows the proportion of the original full LD matrix approximated by the low-dimensional representation, measured in matrix (Frobenius) norm. The fine-tuned hyperparameter settings for different strategies are: (*b, r*) = (500,700) for “Seq_Band_LR”, (*b, r*) = (400,800) for “PSD_Band_LR”, and (*b, r*) = (400,800) for “Joint_Band_LR”, which are selected based on minimizing the CV bias. The runtime reported is for the HEELS heritability estimation.

| Strategy | Average across hyperparameter settings |  |  |  |  |  |
| --- | --- | --- | --- | --- | --- | --- |
|  | % Approx of LD | Heritability estimates |  |  | Runtime |  |
|  |  | Bias | Variance | MSE | In seconds | % saved |
| Exact_LD | 100% | 0.00134 | 0.00001 | 0.00002 | 1578.24 | - |
| Seq_Band_LR | 93.06% | 0.09576 | 0.01808 | 0.06663 | 1428.00 | 9.52% |
| PSD_Band_LR | 92.89% | 0.04135 | 0.00006 | 0.00205 | 905.81 | 42.61% |
| Joint_Band_LR | 94.73% | 0.07530 | 0.00045 | 0.00733 | 830.67 | 47.37% |
| Strategy | Fine-tuned hyperparameter setting |  |  |  |  |  |
|  | % Approx of LD | Heritability estimates |  |  | Runtime |  |
|  |  | Bias | Variance | MSE | In seconds | % saved |
| Seq_Band_LR | 93.97% | 0.00009 | 0.00002 | 0.00002 | 959.38 | 39.21% |
| PSD_Band_LR | 93.84% | 0.01972 | 0.00002 | 0.00041 | 608.42 | 61.45% |
| Joint_Band_LR | 95.96% | 0.04140 | 0.00004 | 0.00175 | 477.79 | 69.73% |

In comparing the performances of the three “Banded + LR” strategies, we observe that they have varying degrees of sensitivity with respect to changes in hyperparameter values *b* and *r* (**Supplementary Figure** 10). Nevertheless, they converge in their approximating abilities (**Supplementary Figure** 11) and lead to small bias when *b* and *r* are sufficiently large (**Figure 2)**. The bandwidth of the banded component plays a critical role in determining the magnitude of the bias in 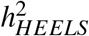, whereas the number of the low-rank factors fine-tunes the estimator (**Supplementary Figure** 12). Among the three “Banded + LR” strategies, we recommend using “PSD_Band_LR” as the default, as it is least sensitive to changes in hyperparameter values, significantly reduces the runtime of HEELS estimation, and produces heritability estimates that are minimally biased (**Supplementary Figure** 10-13).

**Figure 2.**
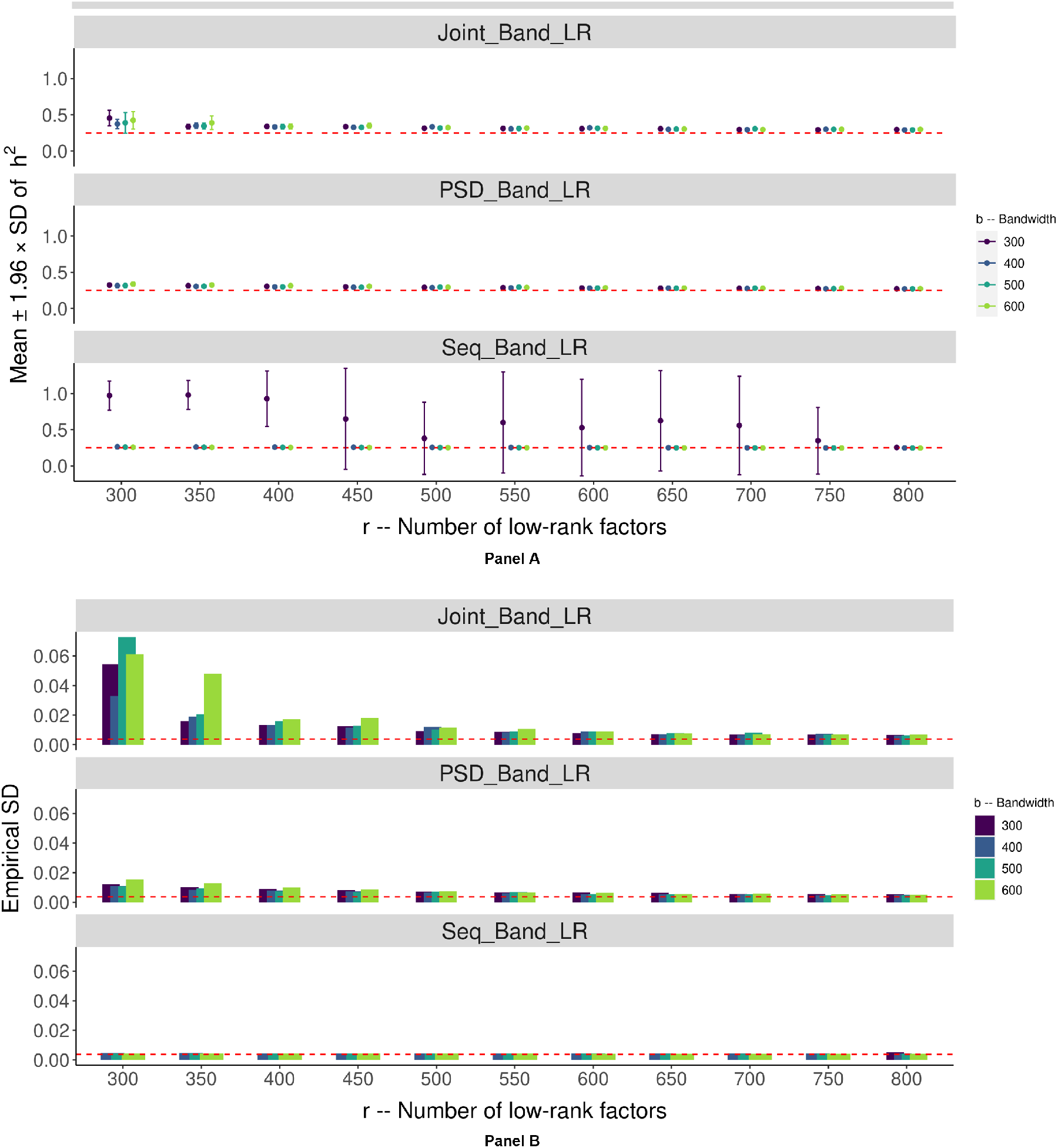
Comparison of the performance of different LD approximation strategies using simulation studies. Simulated phenotypes using real genotypic data from UK Biobank, array SNPs on chromosome 22 with MAF > 0.01 (*n* = 332,430, *p* = 9,220). Three LD approximation methods using the Banded + LR structure with different choices of bandwidths *b* and ranks *r* are compared: Joint_Band_LR, PSD_Band_LR, Seq_Band_LR (**Online Methods** and **Supplemental Notes**). **Panel A**: 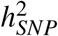 estimates. Red reference line represents the true heritability value of 0.25. The upper (lower) whisker extends from the mean to the values 1.96 × *SE* above (below) the mean. **Panel B**: Empirical SD. Red reference line corresponds to the empirical SD when no approximation is applied (*i.e*. full LD is used).

Finally, we developed a data-adaptive procedure to select the optimal hyperparameter values of *b* and *r* using cross-validation (CV). We refer to this procedure as “pseudo-validation”, as it does not require actual held-out samples but only relies on simulated summary statistics (**Online Methods**). We validated the effectiveness of this procedure, by verifying that the optimal hyperparameter values obtained from this algorithm indeed yield 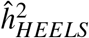 estimates with the highest accuracy and precision (**Supplementary Figure** 14). To improve the computational efficiency of our tuning procedure, we implemented our own version of the incremental SVD algorithm^40^, and adopted several computational techniques to speed up LD decomposition and the tuning algorithm (**Supplementary Notes**).

### Precise estimates of local heritability in the UK Biobank

We applied HEELS to estimate the local SNP-heritability of 30 anthropometric, medical and behavioral traits in the UK Biobank (**Online Methods**), using the LD blocks estimated by Berisa and Pickrell^41^, which have been widely used to proxy approximately independent loci on the human genome^42–44^. We used Z-statistics for all phenotypes and interpret heritability on the liability scale for binary traits^8,45,46^. In line with our expectation based on the theoretical and simulation results, there is an exceptionally high concordance rate between the local heritability estimates from HEELS and those from REML (*r*^2^ = 0.98 on average across the traits), which is the gold standard or the most efficient estimator but requires individual-level data (**Figure 3)**. In contrast, the correlations between the GRE or HESS estimates and the REML estimates are weaker (*r*^2^ = 0.88).

**Figure 3.**
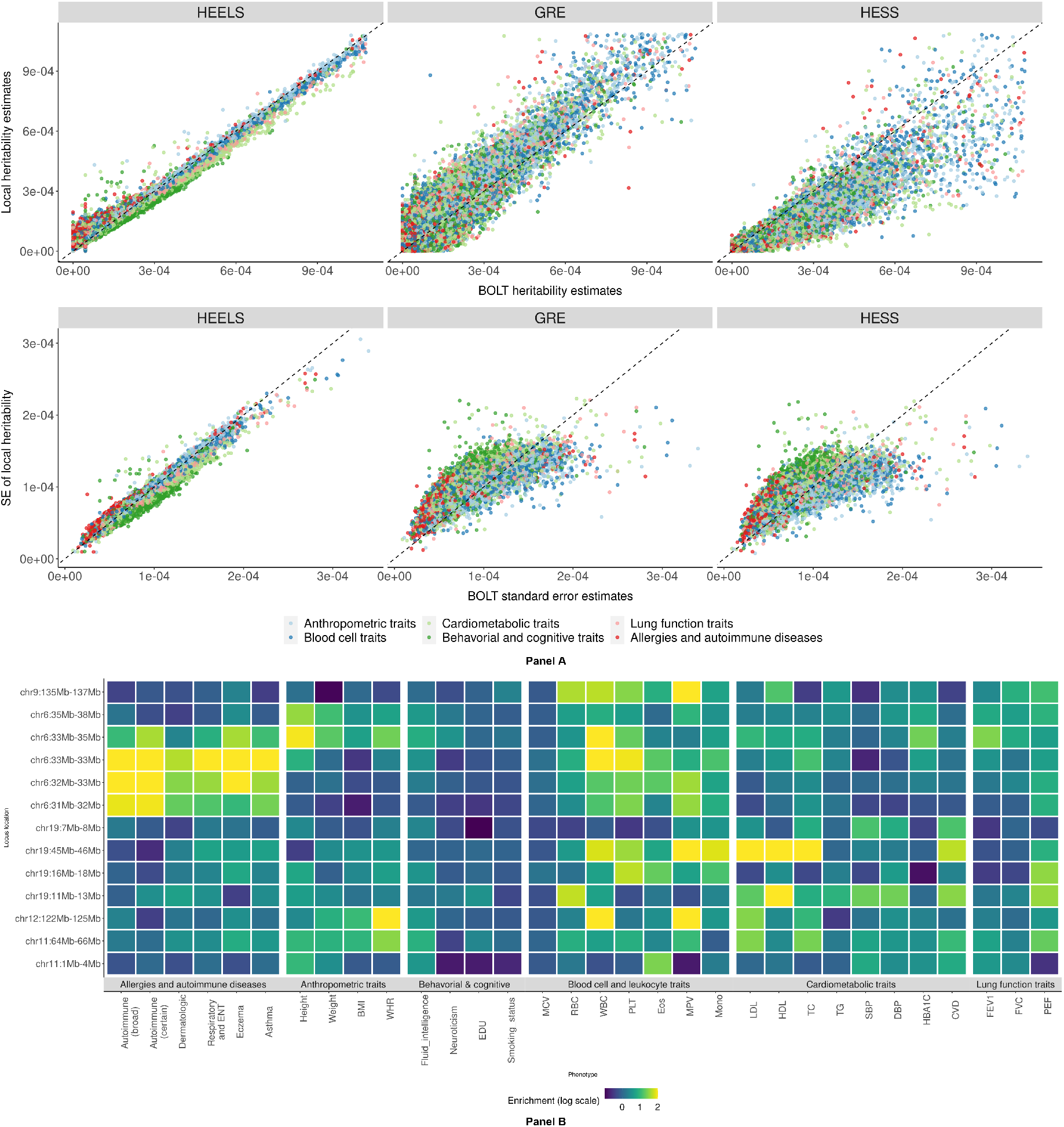
Analysis results from application of HEELS to UKB. **Panel A**: Comparison of heritability estimates and standard errors between BOLT-REML and summary-statistics-based methods (HEELS, GRE, HESS) (*n* = 332,340). Each dot represents a local estimate at one locus for a given trait. Red dashed line: *y* = *x*. **Panel B**: Heritability enrichment of multiple diseases and traits at putative pleiotropic risk loci. Loci locations on the Y-axis are denoted by the start and end positions (in Mbp). The shade of each box represents the enrichment of heritability for the given region and trait, with log transformation.

HEELS yields standard errors that are most comparable to those from individual-level data based REML and its relative efficiency is higher than the other existing summary-statistics based methods (**Supplementary Table** 3). We applied our hyperparameter tuning algorithm to obtain the optimal representations of the LD blocks, and found that these LD approximations generally perform well, yielding local *h*^2^ estimates that are similar to those which are based on using the full LD (**Supplementary Figure** 16). The “Joint_Band_LR” strategy leads to lower accuracy loss compared to the other two strategies (“Seq_Band_LR” and “PSD_Band_LR”), plausibly due to its greater capacity of approximating the LD (**Supplementary Figure** 18).

### Applications of HEELS: Contrasting polygenicity of complex traits and diseases; Identifying risk loci with putative pleiotropic effects

Studies have shown that complex traits and diseases have different degrees of polygenicity^18^. Local heritability estimation provides a useful tool for comparing the polygenic architecture of complex traits, as its distribution pattern across the genome can reflect the degree of which the genetic variance is spread among loci^12,47^. Using local heritability estimated with greater precision from HEELS, we found varying degree of polygenicity across a wide range of traits (**Supplementary Figure** 19). For instance, the genetic variances of behavioral traits and anthropometric traits are more evenly spread out across the genome, whereas those of autoimmune diseases and allergic conditions, lipid traits are more localized and disproportionately dispersed on the genome. Alternatively, we evaluated the polygenicity of complex traits by examining the Pearson correlation coefficients between the genetic variances and the sizes of genomic regions. Indeed, we observed a near-perfect linear relationship between chromosome length and the fraction of 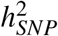 explained by the chromosome for highly polygenic traits, such as BMI (*r*^2^ = 0.988), WHR (*r*^2^ = 0.919), educational attainment (*r*^2^ = 0.991) and neuroticism (*r*^2^ = 0.990). In contrast, the correlations are much lower for less polygenic traits, such as HDL (*r*^2^ = 0.295), LDL (*r*^2^ = 0.537) and autoimmune diseases (*r*^2^ = 0.757) (**Supplementary Figure** 20).

Next, we used the local heritability estimated from HEELS to identify risk loci that are highly enriched for related traits and thus potentially host pleiotropic effects (**Figure 3)**. Our results recapitulate known disease-associated loci from previous findings. For example, we found that the Major Histocompatibility Complex (MHC) (Chr6:31Mb-34Mb) shows strong pleiotropic signals for immunologically relevant diseases and leukocytes involved in innate immunity and inflammatory response. This region, often referred as the human leukocyte antigen (HLA), is known to be highly polygenic^48^ and plays a key role in the induction and regulation of immune responses^49–51^. We also identified Chr19:45Mb-46Mb as a heritability hotspot, which is highly enriched for multiple lipid and cardiometabolic traits while being moderately enriched for leukocytes. This locus harbors the gene cluster – *APOE, APOC1, APOC4, APOC2* – which code for apolipoproteins that are known to be responsible for controlling plasma lipid levels, with subsequent implications in cardiovascular pathology^52^. Our results highlight the probable presence of pleiotropic effects regulating lipid metabolism^53,54^.

We identified two loci that may provide novel insights into the shared genetic basis between related traits and diseases. First, we found that the Chr19:16-18Mb locus is not only enriched for leukocytes such as platelet, lymphocyte and eosinophil, but also enriched for peak expiratory flow. Recent studies have associated platelet-to-lymphocyte ratio (PLR) with the severity of COVID-19, as the PLR of patients can indicate the degree of cytokine storm^55,56^. Our results suggest a putative link between the inflammatory markers and lung functions on a genetic level, and point to a specific region with pleiotropic effects. Further study of this locus may help illustrate its prognostic value for impaired pulmonary functions due to systemic inflammation.

Another region of particular interest is Chr4:143Mb-146 Mb, which is highly enriched for all three of the lung function traits in our analysis (**Supplementary Figure** 17). Although no evidence has directly implicated this region for lung functions, the 4q31 locus – which contains this block – has been prioritized in three GWAS studies^57–59^ and one meta-analysis^60^. One plausible explanation for the pleiotropic effects of this region is that it hosts the Hedgehog (Hh)-interacting protein (*HHIP*) gene, which is associated with multiple pulmonery traits, such as FEV1/FVC ratio, COPD and lung cancer^61–64^. As a regulator of the Hh signalling pathway, *HHIP* is vital for embryonic lung development and is also involved in mature airway epithelial repair^65^. We hypothesize that the observed pleiotropic effects at this locus may be attributable to the *HHIP* gene and its broad influence on lung functionality^66,67^.

## Discussion

To summarize, we have introduced in this paper a REML-based approach HEELS to obtain highly efficient variance component and heritability estimators in linear mixed models, using marginal GWAS association summary statistics and in-sample LD matrix (i.e. empirical covariance matrix of the genotypes). Our estimator only requires summary-level statistics, but yields a highly efficient heritability estimator that is comparable to the REML estimator based on individual-level data, and outperforms the existing summary statistics based heritability methods such as LDSC, GRE and HESS, in terms of statistical efficiency. Another important contribution of our work is that we showcase the benefits of approximating the empirical LD matrix using a banded matrix plus a low-rank matrix. We demonstrate in simulations that this low-dimensional representation can improve the computational efficiency of our algorithm while incurring minimal loss of accuracy or efficiency in estimating 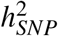. The pseudo-validation procedure we propose provides a principled way to select hyperparameters, in contrast to existing approaches which make arbitrary assumptions about the structure of the empirical LD. We emphasize that our proposed low-dimensional approximations of the LD matrix can be employed more generally to facilitate the storage and sharing of large LD matrices.

We discuss several limitations of our current work and outline some future directions. First of all, our estimator is derived based on the LMM, which assumes a normal distribution of the causal effect sizes. This assumption can greatly simply our comparison between HEELS and REML and help illustrate our main conceptual advance relating to statistical efficiency, but it can be relaxed in two main ways. One is to allow zero effects and incorporate some “sparse” components in the architecture. This direction has been explored extensively in the Bayesian variable selection literature, leading to the development of BSVR^31^, BSLMM^25^ and the Bayesian-alphabet models^68^. The other direction is to adopt a more flexible heritability model, where SNP-specific contribution to the genetic variance can vary. This alternative scheme was put forward by authors of the LDAK^9^ model and examined carefully in the subsequent works. We hope to extend HEELS such that it can accommodate both sparse effects and marker-specific weights, but this extension is challenging, especially because we hope to retain the high statistical efficiency of HEELS.

Second, we emphasize that HEELS requires and relies on the *in-sample* or empirical LD estimated from the GWAS sample. This restriction relates to our definition of heritability in the LMM where we condition on **X**. Our preliminary analyses suggest that replacing the empirical LD with the population or out-of-sample LD in HEELS can lead to biased heritability estimates (**Supplementary Figure** 23). Further research is needed to investigate how out-of-sample LD can influence the statistical properties of the HEELS estimator. HEELS can potentially be applied to summary statistics from meta-analysis summary statistics released by large consortia. The key challenge of applying HEELS to meta-analyzed summary statistics is the appropriate integration of the LD information from multiple cohorts. One fruitful direction for future research is to explore ways to combine the low-dimensional representations of the LD matrices from different studies^69^. We note that in rare variant association tests, researchers routinely release both the vector of score statistics and the covariance (LD) matrix when publishing the results^70–73^. While this is not yet standard practice for common variant analysis, we advocate including the in-sample LD matrix as part of the summary-level statistics when publishing GWAS results. This will enable a more precise characterization of the disease’s genetic architecture (e.g. through HEELS), facilitate privacy-preserving benchmarking efforts^74^, and alleviate challenges that are associated with external LD panels, especially in the context of meta-analyses^75^.

Lastly, we note that the HEELS estimator is best suited for studying and contrasting the heritabilities of *local* genomic regions (up to 20,000 markers). Applying HEELS to a larger set of markers (e.g. genome-wide estimation) can be difficult for two main reasons. First, the computational cost of running HEELS scales quadratically with the number of markers, so the estimation will require additional computational resources when applied to larger blocks on the genome. Second, we determined in simulations that aggregating local heritability estimates can lead to bias in global heritability estimates, a phenomenon that has previously been observed^10^ and may be attributable to LD leakage, *i.e.,* non-zero correlations between SNPs on different blocks^12,38^. To address these challenges, we are actively developing *scalable* algorithms to optimally approximate the LD using a banded + low-rank representations. This advance is not within the scope of our current work, but will make HEELS applicable to estimate the total genetic variance of hundreds of thousands of markers.

## Online Methods

### Statistical model

Let **y** be a length-*n* vector that denotes the phenotypes of *n* samples. Denote by **X** ∈ ℝ^*n*×*p*^ the genotype matrix of *n* individuals based on *p* markers or SNPs. We standardize **X** and **y** such that the variance of the phenotype is 1 and the variance of each marker-specific genotype vector is 1/*p*, or *diag*(**X**^⊤^**X**/*n*) = 1/*p*. Let **S** and **R** denote the the marginal association statistics and the in-sample LD matrix, *i.e*. 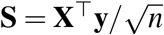 and **R** = **X**^⊤^**X**/*n*. Our goal is to develop a heritability estimator using the two statistics (**S**, **R**), which attains comparable statistical efficiency as the REML estimator based on individual-level data (**X**, **y**). We start by considering the likelihood function, assuming individual-level data can be accessed.

We use an additive genetic model for the phenotypes as **y** = **X***β* + *ε*, where ***β*** is a *p* × 1 vector assumed to follow 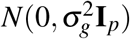, and *ε* is a length-*n* vector distributed as 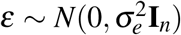. Under these assumptions, **y** ~ *N*(0, **V**), where the variance-covariance matrix is 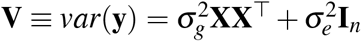. We define SNP-heritability conditional on **X** as the following,

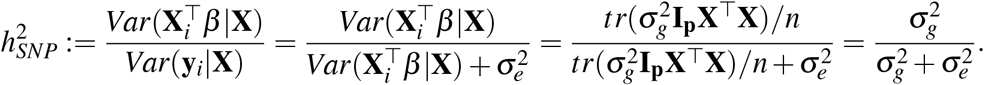

The log-likelihood function for 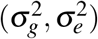 is,

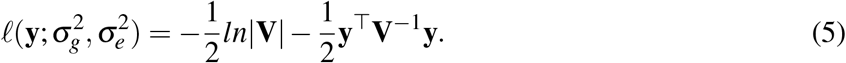

Henderson developed a set of equations^16,76^, known as the mixed model equations (MME), which maximize the joint density of the outcomes and the random effects,

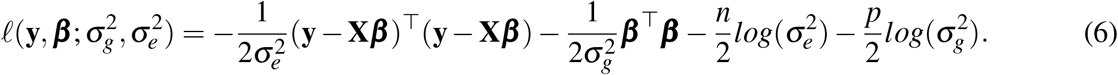

The Best Linear Unbiased Predictor (BLUP), which are estimates for the random effects from these MMEs, can be plugged into the score equations (2)-(3) to generate an iterative procedure for estimating the variance components.^17,77^ We exploit the “dual” form of this algorithm, which gives rise to the HEELS estimator. Note that for simplicity, we have assumed all of the observable environmental factors have been projected out, but covariates (*i.e*. fixed effects) can be easily incorporated into the model by adopting the restricted maximum likelihood approach and using the projection matrix.^78,79^

### The HEELS procedure

HEELS uses the marginal association statistics **S** = **X**^⊤^**y** and the LD matrix **R** = **X**^⊤^**X** to solve for the variance component estimates that maximize the likelihood in (7), alternating between updating the BLUP estimates 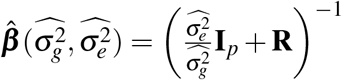 and updating the variance component estimates 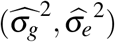 until convergence. The marginal likelihood in (5) can be expressed using the joint likelihood (6) and the probability of the causal effects, using the partition theorem (**Supplementary Notes**),

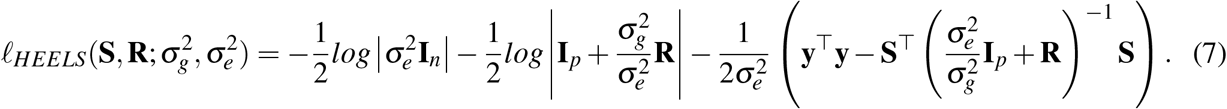

It can be shown that the score equations of 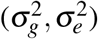 derived from the likelihood in (5) are identical to the score equations based on (7), giving rise to our HEELS updating equations (2) and (3) (**Supplemental notes**). The BLUP estimates can be viewed as ridge estimators of the joint effect sizes, where the penalty coefficient is set to the current value of 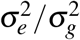 at each iteration. We update 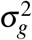 by the mean sum of squares of the BLUP estimates, with the average taken over the effective number of markers, *p* – *tr*([**W**]^−1^) to account for the bias introduced via the ridge penalty.

We emphasize that the HEELS algorithm does not entail any standalone individual-level data **X** or **y**, but only depend on the summary statistics **S**, **R**. The involvement of phenotype variance **y**^⊤^**y** in equation (3) does not necessarily imply a requirement to access **y**, especially if the phenotypes and genotypes are standardized (see similar discussion in Zou *et al*.^23^). In practice, when only the Z-statistics are available, we scale the estimates of 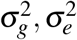 at each iteration as: 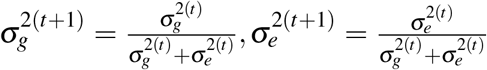, which is equivalent to approximating **y**^⊤^**y**/*n* by 1. We conducted extensive simulations to validate that the HEELS estimator remains unbiased when such an approximation is applied, although the efficiency of our estimator can be slightly inflated. We note that the HEELS procedure can be viewed as equivalent to an EM algorithm (**Supplementary Notes**), where the resulting estimates are stable numerically and belong to the parameter space^80^. This is an important feature that is not universally shared with other REML estimation algorithms such as the Anderson’s algorithm^77,81^ and the Newton-Raphson’s algorithm^82^.

### Standard error of the HEELS estimator

We derive the analytic variance of the HEELS estimator using the Fisher information matrix. For true values of the variance components 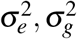 and a positive definite variance-covariance matrix **V**, the Fisher Information Matrix is (**Supplementary Notes**):

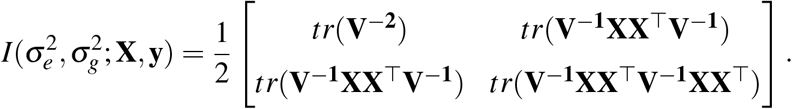

Using properties of trace and the identity of 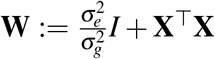, we can rewrite the Fisher information using the summary statistics and the LD matrix as the following^80^:

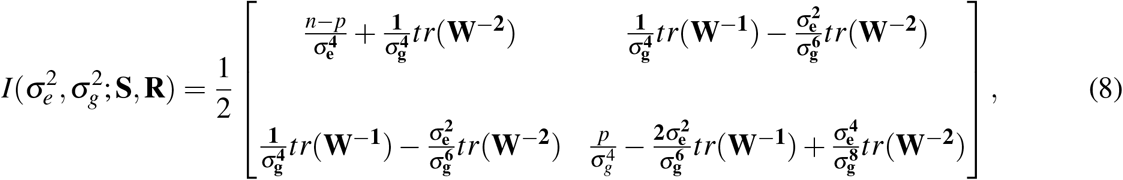

where **W** is defined in the same way as in equation (1). To obtain a variance estimator for 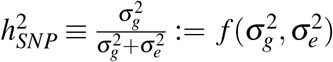, we apply the multivariate Delta method, and use the plug-in estimator:

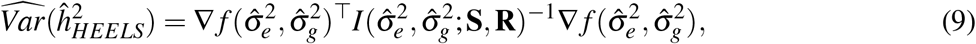

where 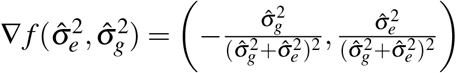. Importantly, (9) only involves **S**, **R** and does not entail **X**, **y**, so we can compute the asymptotic variance of 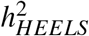 without resorting to individual-level data.

### Low dimensional representations of the LD matrix

Several low-dimensional representations of the LD matrix have been employed in the literature – for example, LD has been approximated by banded matrices^36,37^; LD reference panel has been regularized via the truncated singular value decomposition (TSVD^)12,38^; other methods have adopted block-diagonal low- rank approximations to account for the LD^20,30^. As far as we know, no previous study has systematically compared the performances of different LD approximation strategies. We close this gap by introducing the “Banded + LR” representation of LD, which also provides a unified framework that facilitates comparisons of the existing LD matrix approximation approaches.

We adopt an optimization approach to solve for the best representation of LD with the “Banded + LR” structure, minimizing 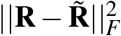, where 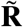 is the working approximating matrix in the form specified in the third column of **Table 2**. For example, the “Joint_Band_LR” approach simultaneously solves for the banded and the low-rank components of the approximation using PSD matrices,

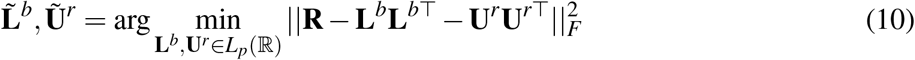

where *L_p_* (ℝ) denotes the set of *p* × *p* lower triangular matrices with real entries. The “PSD_Band_LR” strategy involves a two-step procedure,

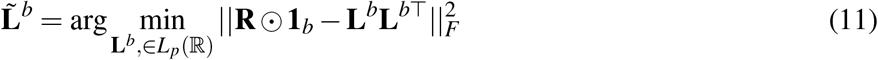

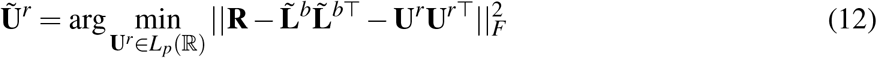

where **1**_*b*_ denotes a square matrix with only the *b* central band equal to 1 and the rest set to 0, and ⊙ is the Hadamard product. In both cases, we approximate **R** as 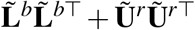.

Below we briefly explain the differences between the low-dimensional representation strategies listed in **Table 2**. When the banded component is a diagonal matrix, the representation is a special case of the “Banded + LR” structure, with *b* = 1, which corresponds to a spiked covariance model^39,83^. We estimate the diagonal elements of the covariance matrix either by taking the smallest eigenvalue of a sub-sample of the LD matrix^84^ (“Spike_LR”), or via joint optimization which solves for the diagonal and the low-rank matrices simultaneously. In the latter case, we allow elements of the diagonal matrix to be identical (“Spike_PSD”) or different (“Spike_PSD_hetero”).

The “Seq_Band_LR” strategy first bands the LD matrix and then performs low-rank decomposition on the residual off-banded matrix. It does not make any PSD assumption on the constituents of its solution. The main advantage of this strategy is that it most accurately represents the banded structure of the LD matrix, *i.e*. the banded component matches exactly the central band of the original LD matrix. A drawback of this approach is its lack of flexibility in approximating the off-central-band structure.

In contrast, the “Joint_Band_LR” strategy jointly approximates the banded and low-rank components using PSD matrices. It provides greater flexibility of approximation for all elements indiscriminately, as the elements of the banded matrix and the low-rank matrix are jointly optimized to minimize the Frobenius norm of the error matrix. A benefit of “Joint_Band_LR” over “Seq_Band_LR” is that due to its explicit minimization of the approximation error, its solution is, by definition, closer to the original LD matrix. A potential shortcoming of the “Joint_Band_LR” strategy, however, is that by imposing a PSD assumption on *both* the banded and the low-rank components of the LD representation, we restrict the solution to a particular set of PSD matrices and it is computationally slower.

To strike a balance between accurately representing the central band of the LD matrix and ensuring the computational efficiency of our algorithm, we developed a hybrid strategy, “PSD_Band_LR”, which first approximates the banded component of the LD matrix using a PSD matrix followed by low-rank decomposing the residual matrix. On one hand, it differs from “Seq_Band_LR” in that the banded component of the approximation is guaranteed to be PSD. On the other hand, “PSD_Band_LR” differs from “Joint_Band _LR” in that it *sequentially* solves for the banded and low-rank constituents of the representation, as opposed to simultaneously. As a result, “PSD_Band_LR” preserves the original structure of the LD matrix better than “Joint_Band_LR” does, leading to a more accurate representation of the central band. “PSD_Band_LR” also exploits the PSD assumption, making it more computationally stable than “Seq_Band_LR” (**Supplementary Notes**).

### Comparison of different LD approximations

The existing LD approximation methods impose different assumptions on the structure of the LD matrix, and the motivation to induce the low-dimensional LD representation varies from case to case. We introduce a unified framework for contrasting the different LD approximation approaches in the context of heritability estimation. For a given LD representation – characterized by its structure (“Strategy”) and the values of its hyper-parameters: the band size *b* and the rank of the low rank component *r* – we considered two measures to evaluate its performance: one is the LD approximation accuracy, measured by the ratio of the Frobenius-norms between the approximation matrix and the target matrix, 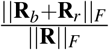; the other is the cross-validation (CV) bias in 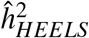, estimated using synthetic phenotypes generated from real in-sample genotypes. The first measure assesses the goodness of the approximation in general, irrespective of the genetic architecture of a trait, whereas the second measure evaluates the approximation performance in the context of heritability estimation.

For low-dimensional representations with smaller *b* and *r*, the performance of the HEELS estimator is greatly influenced by the assumed structure of the approximation strategy and the hyperparameter values. For example, for each of the “Banded + LR” strategies, we could pinpoint the approximate range of an underlying transition point (*b_min_*), which marks the minimum optimal value of *b* – widening the bandwidth up to this value can substantially reduce the bias in 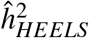, but further increasing *b* beyond this threshold value has a diminishing “de-biasing” effect on 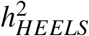 (**Supplementary Figure** 12).

Banding the full LD first and then approximating the residual using a low-rank matrix (“Seq_Band_LR”) yields the most unbiased heritability estimates when *b* and *r* are sufficiently large, but can produce estimates with large bias if *b* and *r* are not appropriately chosen; approximating the LD matrix by simultaneously solving for the banded and low-rank components (“Joint_Band_LR”) leads to less-biased estimates even when *b* and *r* are sub-optimal, although solving for the approximation is more computationally expensive than “Seq_Band_LR”; the hybrid strategy which approximates the banded component using a PSD matrix first and finds the low-rank decomposition of the residual (“PSD_Band_LR”) is least sensitive to changes in hyperparameter values and produces heritability estimates that are most stable across LD approximation settings.

### Algorithms for hyperparameter tuning

An important aspect of our low-dimensional LD representation algorithm is the tuning of the hyper-parameters. While heuristics or prior knowledge about the structure of the LD can be used to determine the optimal values of (*b, r*), we used a more principled way to evaluate the performance of low-dimensional LD representation. We propose using a data-adaptive procedure to identify the best low-dimensional representation of the LD matrix, using synthetic phenotypic data and cross-validation (**Supplementary Notes**).

To facilitate the searching of the optimal number of low-rank factors, *r*\*, we implemented our own version of the “incremental SVD algorithm”, where the number of low-rank factors increases step by step and the performance of the HEELS estimator is evaluated dynamically. To speed up the low-rank decomposition, we replaced the *exact* solutions based on direct eigen-decomposition with the *approximate* solutions, and compared two approaches – one is the optimization approach (“optim”), where we solve 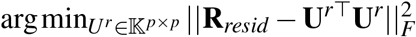, and **R**_*resid*_ is the residual off-banded component to be decomposed or approximated and 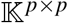 is the collection of lower triangular matrices; the other approach is based on random sketching (“random”), where we applied fast PCA to decompose ***R***_*resid*_ via SVD^84^. The simulation results indicate that using the optimization approach is less prone to bias and is more robust to variation in the hyperparameter values (**Supplementary Figure** 15). While we observe larger bias when the random sketching SVD approach is used to approximate the low-rank component, it can be more advantageous in applications with larger problem size.

### Simulation framework

We obtained 332, 340 unrelated White British individuals by extracting samples with self-reported British ancestry who are more than third-degree relatives and excluding subjects with putative sex chromosome aneuploidy. We used genotyped array SNPs only to ensure high quality of measurement, and filtered out variants with genotype missingness > 0.01 or have MAF greater than 0.01. For experiments that involve subsets of the UKB individuals, we recalculated minor allele frequency and re-applied the filters above to the subsample. The environmental effects are drawn independently from the genetic effects for each individual. This ensures that population structure or cryptic relatedness among individuals will have minimal impact on our estimates in the simulations.

We simulated phenotypes with different genetic architectures using real genotypes from the UK Biobank, varying sample size (*n*), the ratio of *n* to the number of markers (*p*) and the degree of polygenicity (*p_causal_*/*p*). Given the raw genotype matrix **G**, we first standardized the genotype matrix: for each SNP *j* and individual *i*, we generated 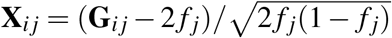, where **G**_*ij*_ ∈ {0,1,2} and *f_j_* is the in-sample minor allele frequency of SNP *j*. For a given degree of polygenicity, we randomly sampled *p_causal_* markers. Denote the set of causal markers by *C*. We drew standardized effect sizes from the distribution, 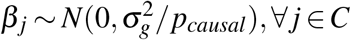, and simulated the phenotype of the i-th individual using *y_i_* = ∑*j*∈*C X_ij_β_j_* + *ε_i_*, where 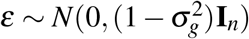. For each genetic architecture, we generated 100 phenotype replicates and obtained 100 estimates using each of the methods we included in the benchmark. Given phenotypes **y** = (*y*_1_,...,*y_N_*)^⊤^ and genotypes 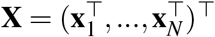, we computed the marginal association statistics using OLS: 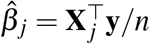. Unless otherwise specified, we used the exact in-sample LD without approximation in the HEELS estimation.

We applied two software to compute the REML estimates. We used GCTA when sample size is small and used BOLT-REML when *n* exceeds 30,000. For experiments with varying sample size, we selected individuals randomly from the full QC-ed UK Biobank data. To create an imbalance of sample size across causal markers, we ranked the marker-specific sample size and defined the lowest 10 percentile of the SNPs as “low-N” SNPs. It is important to note that we did not include any markers that would have been filtered out by conventional QC steps, such as call rate of genotypes > 0.90, *i.e*. define low-sample size relatively and dynamically *within* the set of high-quality SNPs that have passed QCs, with varying sample size across different settings.

Our primary metric of interest is relative efficiency, which is the ratio of the variance of the REML estimator and the variance of a given heritability estimator. We also compared the bias and the mean squared error of the estimators across different simulation settings.

### UKB empirical analysis

We applied our method to the unrelated white British individuals in the UK Biobank, using common variants with MAF > 1% on the UK Biobank Axiom array (*n* = 332,430, *p* = 533,169). We estimated the local SNP-heritability of 33 phenotypes from different trait domains, including anthropometric traits (height, weight, BMI and waist-to-hip ratio), hematological traits (mean corpuscular volume, red blood cell count, white blood cell count, platelet count, eosinophils, lymphocytes, mean platelet volume and monocytes), lipid or metabolic traits (LDL, HDL, triglycerides, total cholesterol), lung function traits (forced expiratory volume, forced vital capacity, peak expiratory flow rate), behavioral traits (smoking behavior, educational attainment, neuroticism) and immunologically relevant traits (autoimmune conditions, asthma, eczema and dermatologic diseases).

We used PLINK to exclude SNPs with MAF <1%, genotype missingness >1%, genotyping rates < 90%, and Hardy-Weinberg disequilibrium p-value > 1e-6. We generated the genome-wide association statistics using the —linear association analyses in PLINK, controlling for age, sex, and the top 40 genetic principal components provided by the UKB^85^. We used BOLT-REML to compute the REML estimates in the UK Biobank, and used the HESS software to compute both the HESS estimates (default setting) and the GRE estimates (*i.e*. no regularization of the in-sample LD matrix), after verifying that the estimates from the two methods indeed coincide when the number of eigenvectors used in TSVD equals to the rank of the genotype matrix. We used the approximately independent loci estimated by Berisa and Pickrell^41^ to define the regions as units of analysis (1.6 Mb on average or 300-400 markers per block using the genotype array data). The 1,703 non-overlapping regions have been been widely used in previous studies^42–44^ and have been demonstrated to capture the true correlation structures among genotypes reasonably well^86^.

## Data availability

This work used genotyped and phenotypes from the UK Biobank study (https://www.ukbiobank.ac.uk) and our access was approved under application number 52008.

## Code availability

Our method (HEELS) has been implemented as an open-source Python package, available at: https://github.com/huilisabrina/HEELS. The following software packages were also used in simulation studies and real data analyses: GCTA (https://yanglab.westlake.edu.cn/software/gcta/#Download); LD Score regression (https://github.com/bulik/ldsc); GRE (https://github.com/bogdanlab/h2-GRE); GRE (https://github.com/bogdanlab/h2-GRE); HESS (https://github.com/huwenboshi/hess); BOLT-LMM v2.3.4 (https://data.broadinstitute.org/alkesgroup/BOLT-LMM/). We also used Python (3.6.3) to perform statistical analyses and used R (3.6.3) for data visualization.

## Acknowledgements

We are very grateful to Luke O’Connor, Huwenbo Shi, Alkes Price, Samuel Kou, Ben Neale and Shamil Sunyaev for their helpful discussions and feedback. This work was supported by grants R35-CA197449, U19-CA203654, R01-HL163560, U01-HG012064, and U01-HG009088 (X. Lin).

## Author contributions statement

H.L., R.M. and X.L. conceived and designed the experiments. H.L. performed the experiments and the statistical analyses. H.L. wrote the manuscript with the participation of R.M. and X.L.

## Notes

### Competing Interest Statement

X. Lin is a consultant of AbbVie Pharmaceuticals and Verily Life Sciences.

### Summary of Updates

Updated introduction, discussion and the relevant sections in the supplementary notes to incorporate more discussion on the relationship between current work and other recently published works.

