## Supplementary Notes for "Accurate and Efficient Estimation of Local Heritability using Summary Statistics and LD Matrix"

#### 1 Literature review of existing SNP-heritability estimation methods

Here we provide a more comprehensive comparison of the existing heritability estimation methods (summarized in **Supplementary Table** 1). It is not our intention to compare HEELS with all or most of the existing heritability estimation methods, but we hope to highlight the similarities and differences among approaches in terms of statistical efficiency.

##### 1.1 Existing heritability estimation methods using individual-level data

Linear mixed models (LMMs) has been widely used as the analytical framework for $h^2_{SNP}$ estimation. In a nutshell, $h^2_{SNP}$ is embedded in the parameters of a LMM as the variance of causal genetic effects. One of the most commonly used variance component estimation methods is restricted maximum likelihood (REML). The well-known tool of Genome-wide Complex Traits Analysis (GCTA) applies this method in the context of genetics studies, which led to the "GREML" estimator of SNP-heritability.[1] GREML uses second-order methods (average information or Fisher scoring) to maximize the log-likelihood function based on individual genotypic and phenotypic data. The solutions are found through iterative optimization algorithms, which can be computationally intensive when applied to large samples.

To overcome the computational limitations GREML, Loh *et al.* introduced BOLT-REML, which is a highly efficient algorithm that enables robust and fast variance component estimation on biobank-scale datasets[2]. BOLT-REML greatly improves the computational efficiency of the estimation procedure of GREML, by approximating the gradient (and the Hessian matrix) of the likelihood function via Monte Carlo sampling. It also adopts numerical techniques such as the conjugate gradient algorithm for mixed-model solutions to reduce its computational cost[2].

Speed *et al.* made an important advance in heritability estimation, by introducing a new REML estimator of $h^2_{SNP}$, called LD-adjusted kinships (LDAK).[3] LDAK differs from GREML in its assumption of the relationships between the effect sizes of causal markers and their minor allele frequencies (MAF) as well as LD tagging. The main advantage of REML estimators, such as GREML and LDAK, is that they have high statistical efficiency – because they produce MLE or REML estimators, especially in large samples, the asymptotic variance of these two estimators is guaranteed to attain the Cramer-Rao lower bound. Previous works have shown that the stratified or multi-component GREML can effectively account for the coupling relationship between causal effect sizes and MAF/LD structure[4]. Therefore, the GREML estimator can be reconciled with the LDAK estimator under certain assumptions[5].

Haseman–Elston (HE) regression is another well-known approach for estimating heritability,[6,7] which is germane to the method-of-moments (MoM) estimators of $h^2_{SNP}$, *i.e.*, population covariance of the phenotypes is matched to the empirical covariance[8]. The computational bottleneck of applying the HE-regression is its requirement to calculate a large genetic relatedness matrix that summarizes the relationship between all $n \times n$ pairs of individuals in the sample, especially when sample size exceeds 30,000[1]. Wu et al. developed a randomized HE-regression (RHE-reg) estimator that reduces both the runtime and the storage requirement of heritability estimation based on HE-regression[9]. The main contribution of their approach pertains to the usage of random sampling for trace calculation and its application of the Mailman algorithm for matrix-vector multiplication to solve the MoM normal equations. Both BOLT-REML and RHE-reg employ random sampling to circumvent frequent and expensive operations that involve high-dimensional matrices. The downside of such approximation is that they tend to produce less statistically efficient estimators, due to the additional variance introduced by randomization[2,9].

Another important line of research estimates SNP-heritability under the Bayesian framework, which adopts different assumptions about the distribution of true causal effect sizes. The first method of this kind is the sparse regression model proposed by Guan and Stephens[10]. They introduced "Bayesian variable selection regression (BVSR)" for SNP-heritability estimation and phenotypic prediction. BVSR assumes that the joint effect size follows a point-normal distribution, and it works well when the genetic architecture of a trait is truly sparse. Later, Bayesian Sparse Linear Mixed Model (BSLMM) was proposed as a hybrid approach that combines the advantages of LMM and BVSR[11]. BSLMM uses a mixture of two normals to jointly model a small number of large effects ("sparse" component) and a large number of small effects ("polygenic" component). A key advantage of the Bayesian framework is its flexibility to model different effect size distributions. For example, the Bayesian alphabet[12] models is a class of variants to BSLMM and BSVR which make different distributional assumptions on the effect sizes, with the BayesC$\pi$ being the closest to BSLMM[13]. The main challenge of using these Bayesian methods however, is that the posterior inference procedure can be quite computationally intensive, *e.g.*, orders of magnitude slower than LMM when applied to large-scale genetic data[11].

##### 1.2 Existing heritability estimation methods using summary statistics

When only GWAS summary-level statistics are available, LD score regression (LDSC) is a state-of-the-art method that is highly computationally efficient and yields heritability estimates that can adjust for confounding by environmental effects or population stratification[14,15]. From the modeling perspective, LDSC is similar to GREML as it also adopts the LMM and assumes that all causal markers contribute to heritability equally. From the estimation perspective, the LDSC estimator of heritability is akin to RHE-reg, as both methods model the second moments of the effect size without making any distributional assumptions. The most important contribution of LDSC is its ability to distinguish confounding bias from true polygenic genetic signals, although a recent study raised some concern about this capacity[16]. SumHer[17] was introduced as a summary-statistics-based extension of LDAK, as it explicitly accounts for the MAF/LD-dependent structure of effect sizes. The comparison between LDSC and SumHer is parallel to the comparison between GREML and LDAK, and recent works have shown the converging performance of these two lines of research with the usage of stratification[4,5,18].

Although summary-statistics-based methods are attractive due to its broader applicability and lessened privacy concern, an important drawback of these approaches is that they generally produce estimates with considerably large standard errors than individual-level data base REML estimates. For example, Zhou *et al.* demonstrated within the framework of Minimal Norm Quadratic Unbiased Estimation (MINQUE) that the weight matrix used by LDSC in solving the MoM normal equations is not optimal, and therefore its estimator does not achieve the highest statistical efficiency[19]. We show the statistical efficiency of our HEELS by establishing its equivalence with the generalized methods-of-moment estimator with the optimal weights under the MINQUE framework. A new MoM estimator – MinQue for Summary Statistics (MQS) – was introduced under this unified framework[19]. The paper shows that MQS can improve the statistical efficiency of LDSC under certain settings, which is affected by the weighting scheme, true heritability as well as the degree of relatedness in the sample.[19] MQS is unbiased and is generally more statistically efficient than LDSC, but still has larger variance than REML (see Figure 2 and Table 2 of Zhou, 2017).

Two other $h^2_{SNP}$ estimators resemble HEELS and are based on summary-level statistics – Generalized Random Estimator (GRE[20]) and Heritability Estimator from Summary Statistics (HESS[21]). Both of these two methods produce closed-form solutions of $h^2_{SNP}$, and are robust to the unknown underlying genetic architecture of the phenotype. Although HESS was developed using a fixed-effect model whereas GRE assumes the causal effect sizes are random, the derivation of these two estimators are closely related, as is evident from their analytical expressions (see later sections). When applied to a local genetic region, $h^2_{HESS}$ coincides with $h^2_{GRE}$ when the in-sample LD is used without any regularization (*i.e.* the number of top eigenvectors used by HESS equals to the rank of the LD matrix).

The summary-statistics-based extension of the Bayesian methods such as BVSR and BSLMM is Regression with Summary Statistics (RSS).[22] RSS adopts similar assumptions as BSLMM but uses the likelihood of the joint effect sizes and performs inference on the SNP-heritability defined using summary statistics. RSS also leverages the banded structure of the LD and uses the shrinkage estimator from Wen and Stephens[23] to approximate LD. The simulation results suggest that RSS is unbiased but is less statistically efficient than its corresponding individual-level data based method (see Figure 3 of Zhu and Stephens, 2017).

A recently developed method, High-Definition Likelihood ("HDL"), shares our goal of improving the statistical efficiency of genetic variance estimators[24]. Although HDL was proposed as a method for genetic correlation estimation, it necessarily computes heritability as an intermediate step. In simulations, we observed bias in HDL's estimates of heritability, and another study has reported similar issues in their benchmarking results[25]. Our approach differs from HDL in two important ways. First, our estimator was derived using a different likelihood function (*i.e.*, we start with the likelihood that assumes individual-level data is known, and then transform the score-solving algorithm into a summary-statistics-based estimating procedure). HDL first derived the likelihood of the marginal statistics, which is closely related to the "RSS" likelihood proposed in Zhu and Stephens[26]. Second, HDL approximates the LD matrix using a combination of banding, blocking and truncated SVD, whereas we proposed approximating the LD matrix in a principled manner using a banded + low-rank representation.

We summarize the SNP-heritability estimator methods reviewed above in **Supplementary Table 1**. We select GREML, LDSC, GRE and HESS as the representative heritability estimation methods to be compared with our HEELS, because the differences between these methods and the other existing approaches lie elsewhere from *statistical efficiency*. For methods that we do not directly compare with HEELS, we cite existing evidence of the statistical efficiency (see the last column of **Supplementary Table** 1). We do not include HDL in our comparisons of summary-statistics-based estimators due to the bias we observed and reported elsewhere, as mentioned above.

#### 2 Model and REML likelihood based on individual-level data

Let $\mathbf{y}$ be a length-$n$ vector that denotes the phenotypes of $n$ samples. Denote by $\mathbf{X} \in \mathbb{R}^{n\times p}$ the genotype matrix of $n$ individuals based on $p$ markers or SNPs. We standardize $\mathbf{X}$ and $\mathbf{y}$ such that the variance of the phenotype is 1 and the variance of each marker-specific genotype vector is $1/p$, or $diag(\mathbf{X}^\top \mathbf{X}/n) = 1/p$. Let $\mathbf{S}$ and $\mathbf{R}$ denote the the marginal association statistics and the in-sample LD matrix, *i.e.* $\mathbf{S} = \mathbf{X}^\top \mathbf{y}/\sqrt{n}$ and $\mathbf{R} = \mathbf{X}^\top \mathbf{X}/n$. Our goal is to develop a heritability estimator using the two statistics $(\mathbf{S}, \mathbf{R})$, which attains comparable statistical efficiency as the REML estimator based on individual-level data $(\mathbf{X}, \mathbf{y})$. We start by considering the likelihood function, assuming individual-level data can be accessed.

We use an additive genetic model for the phenotypes as $\mathbf{y} = \mathbf{X}\beta + \varepsilon$, where $\boldsymbol{\beta}$ is a $p \times 1$ vector assumed to follow $N(0, \sigma_g^2 \mathbf{I}_p)$, and $\varepsilon$ is a length-$n$ vector distributed as $\varepsilon \sim N(0, \sigma_e^2 \mathbf{I}_n)$. Under these assumptions, $\mathbf{y} \sim N(0, \mathbf{V})$, where the variance-covariance matrix is $\mathbf{V} \equiv var(\mathbf{y}) = \sigma_g^2 \mathbf{X}\mathbf{X}^\top + \sigma_e^2 \mathbf{I}_n$. We define SNP-heritability conditional on $\mathbf{X}$ as the following,

$$h^2_{SNP} := \frac{Var(\mathbf{X}_i^\top \beta|\mathbf{X})}{Var(\mathbf{y}_i|\mathbf{X})} = \frac{Var(\mathbf{X}_i^\top \beta|\mathbf{X})}{Var(\mathbf{X}_i^\top \beta|\mathbf{X}) + \sigma_e^2} = \frac{tr(\sigma_g^2 \mathbf{I_p} \mathbf{X}^\top \mathbf{X})/n}{tr(\sigma_g^2 \mathbf{I_p} \mathbf{X}^\top \mathbf{X})/n + \sigma_e^2} = \frac{\sigma_g^2}{\sigma_g^2 + \sigma_e^2}.$$

The log-likelihood function for $(\sigma_g^2, \sigma_e^2)$ is,

$$\ell(\mathbf{y}; \sigma_g^2, \sigma_e^2) = -\frac{1}{2} ln|\mathbf{V}| - \frac{1}{2} \mathbf{y}^\top \mathbf{V}^{-1} \mathbf{y}. \quad (1)$$

Using well-known results in matrix differentiation, we can maximize this log-likelihood with respect to $\sigma_g^2$ and $\sigma_e^2$, by solving the following score equations[27]:

$$U_{\sigma_g^2}(\mathbf{y}) = -\frac{1}{2} tr(\mathbf{X}^\top \mathbf{V}^{-\mathbf{1}} \mathbf{X}) + \frac{1}{2} \mathbf{y}^\top \mathbf{V}^{-\mathbf{1}} \mathbf{X}\mathbf{X}^\top \mathbf{V}^{-\mathbf{1}} \mathbf{y} = \mathbf{0} \quad (2)$$

$$U_{\sigma_e^2}(\mathbf{y}) = -\frac{1}{2} tr(\mathbf{V}^{-1}) + \frac{1}{2} \mathbf{y}^\top \mathbf{V}^{-\mathbf{1}} \mathbf{V}^{-\mathbf{1}} \mathbf{y} = \mathbf{0} \quad (3)$$

Henderson developed a set of equations[28,29], known as the mixed model equations (MME), which maximize the joint density of the outcomes and the random effects,

$$\ell(\mathbf{y}, \beta; \sigma_g^2, \sigma_e^2) = -\frac{1}{2\sigma_e^2} (\mathbf{y} - \mathbf{X}\boldsymbol{\beta})^\top (\mathbf{y} - \mathbf{X}\boldsymbol{\beta}) - \frac{1}{2\sigma_g^2} \boldsymbol{\beta}^\top \boldsymbol{\beta} - \frac{n}{2} log(\sigma_e^2) - \frac{p}{2} log(\sigma_g^2). \quad (4)$$

The Best Linear Unbiased Predictor (BLUP), which are estimates for the random effects from these

MMEs, can be plugged into the score equations (2)-(3) to generate an iterative procedure for estimating the variance components.[27,30] We exploit the "dual" form of this algorithm, which gives rise to the HEELS estimator. Note that for simplicity, we have assumed all of the observable environmental factors have been projected out, but covariates (*i.e.* fixed effects) can be easily incorporated into the model by adopting the restricted maximum likelihood approach and using the projection matrix.[31,32]

### 3 HEELS updating equations

The marginal likelihood in (1) can be expressed using the joint likelihood (4) and the probability of the causal effects, using the partition theorem,

$$\begin{aligned} f(\mathbf{y}) &= \int_{\boldsymbol\beta} f(\mathbf{y},\boldsymbol\beta)d\boldsymbol\beta = \int_{\boldsymbol\beta} f(\mathbf{y}|\boldsymbol\beta)f(\boldsymbol\beta)d\boldsymbol\beta \\ &= \int_{\boldsymbol\beta}(2\pi\sigma_g^2)^{-p/2}(2\pi\sigma_e^2)^{-n/2}exp\left(-\frac{1}{2}\left(\frac{1}{\sigma_e^2}(\mathbf{y}-\mathbf{X}\boldsymbol\beta)^\top(\mathbf{y}-\mathbf{X}\boldsymbol\beta)+\frac{1}{\sigma_g^2}\boldsymbol\beta^\top\boldsymbol\beta\right)\right)d\boldsymbol\beta \\ &= \left|\mathbf{X}^\top\mathbf{X}+\frac{\sigma_e^2}{\sigma_g^2}\mathbf{I}\right|^{p/2}\left(2\pi\sigma_e^2/\sigma_g^2\right)^{p/2}(2\pi\sigma_e^2)^{-n/2}exp\left(-\frac{1}{2\sigma_e^2}\left(\mathbf{y}^\top\mathbf{y}-\mathbf{y}^\top\mathbf{X}\left(\mathbf{X}^\top\mathbf{X}+\frac{\sigma_e^2}{\sigma_g^2}\mathbf{I}\right)^{-1}\mathbf{X}^\top\mathbf{y}\right)\right). \\ &\qquad \text{(use the kernel of } \boldsymbol\beta\sim N\left(\left(\mathbf{X}^\top\mathbf{X}+\tfrac{\sigma_e^2}{\sigma_g^2}\mathbf{I}\right)^{-1}\mathbf{X}^\top\mathbf{y},\tfrac{1}{\sigma_e^2}\mathbf{X}^\top\mathbf{X}+\tfrac{1}{\sigma_g^2}\mathbf{I}\right)\text{)} \end{aligned}$$

Therefore, taking the log and omitting the constant term, we can rewrite equation (1) as a function of summary statistics, given the unknown variance components,

$$\ell_{HEELS}(\mathbf{S},\mathbf{R};\sigma_g^2,\sigma_e^2) = -\frac{1}{2}log\left|\sigma_e^2\mathbf{I}_n\right| - \frac{1}{2}log\left|\mathbf{I}_p+\frac{\sigma_g^2}{\sigma_e^2}\mathbf{R}\right| - \frac{1}{2\sigma_e^2}\left(\mathbf{y}^\top\mathbf{y}-\mathbf{S}^\top\left(\frac{\sigma_e^2}{\sigma_g^2}\mathbf{I}_p+\mathbf{R}\right)^{-1}\mathbf{S}\right). \quad (5)$$

HEELS uses the marginal association statistics $\mathbf{S}=\mathbf{X}^\top\mathbf{y}$ and the LD matrix $\mathbf{R}=\mathbf{X}^\top\mathbf{X}$ to solve for the variance component estimates that maximize the likelihood in (5), alternating between updating the BLUP estimates $\hat{\boldsymbol\beta}(\widehat{\sigma_g^2},\widehat{\sigma_e^2}) = \left(\frac{\widehat{\sigma_e^2}}{\widehat{\sigma_g^2}}\mathbf{I}_p+\mathbf{R}\right)^{-1}\mathbf{S}$ and updating the variance component estimates $(\widehat{\sigma_g}^2,\widehat{\sigma_e}^2)$ until convergence. Below we provide the full details for deriving the updating equations used in HEELS, for which we referenced Chapter 7 of Searle, Casella and McCulloch[33] as well as Harville (1977).[27]

#### 3.1 Updating equation for $\beta$

The BLUP estimates that maximize the joint density in equation (4) satisfy $\left(\frac{\mathbf{R}}{\sigma_e^2}+\frac{1}{\sigma_g^2}\right)\boldsymbol\beta = \frac{1}{\sigma_e^2}\mathbf{S}$. Therefore, if the variance components are given and known, we can calculate the BLUP estimates as $\boldsymbol\beta =$

$\left(\frac{\sigma_e^2}{\sigma_g^2}\mathbf{I}_p + \mathbf{R}\right)^{-1}\mathbf{S}$. This is essentially our updating equation for the joint effect sizes if we replace the parameters with their current estimates at each iteration.

#### 3.2 Updating equation for $\sigma_e^2$

Define $\mathbf{H} = \frac{\mathbf{V}}{\sigma_e^2}$, so that $\mathbf{V}^{-1} = \frac{\mathbf{H}^{-1}}{\sigma_e^2}$. We can then establish the following:

$$\mathbf{X}\mathbf{X}^\top \sigma_g^2 = \mathbf{V} - \sigma_e^2 \mathbf{I} = (\mathbf{H} - \mathbf{I})\sigma_e^2.$$

Setting $U_{\sigma_g^2} = \mathbf{0}$, we have:

$$\begin{aligned}
tr(\mathbf{X}^\top \mathbf{V}^{-1}\mathbf{X}) &= (\mathbf{X}^\top \mathbf{V}^{-1}\mathbf{y})^\top (\mathbf{X}^\top \mathbf{V}^{-1}\mathbf{y}) \\
tr(\mathbf{V}^{-1}\mathbf{X}\mathbf{X}^\top \sigma_g^2) &= \mathbf{y}^\top \mathbf{V}^{-1}\mathbf{X}\mathbf{X}^\top \sigma_g^2 \mathbf{V}^{-1}\mathbf{y} \\
tr(\mathbf{V}^{-1}\mathbf{X}\mathbf{X}^\top \sigma_g^2) &= \mathbf{y}^\top \mathbf{V}^{-1}(\mathbf{V} - \sigma_e^2\mathbf{I})\mathbf{V}^{-1}\mathbf{y} \\
tr(\mathbf{V}^{-1}\mathbf{X}\mathbf{X}^\top \sigma_g^2) &= \mathbf{y}^\top \mathbf{V}^{-1}(\mathbf{V} - \sigma_e^2\mathbf{I})\mathbf{V}^{-1}\mathbf{y} \\
tr(\mathbf{V}^{-1}\mathbf{X}\mathbf{X}^\top \sigma_g^2) &= \mathbf{y}^\top \mathbf{V}^{-1}(\mathbf{H} - \mathbf{I})\sigma_e^2 \mathbf{V}^{-1}\mathbf{y} \\
tr\left(\frac{\mathbf{H}^{-1}}{\sigma_e^2}(\mathbf{H} - \mathbf{I})\sigma_e^2\right) &= \mathbf{y}^\top \frac{\mathbf{H}^{-1}}{\sigma_e^2}(\mathbf{H} - \mathbf{I})\sigma_e^2 \frac{\mathbf{H}^{-1}}{\sigma_e^2}\mathbf{y} \\
tr(\mathbf{H}^{-1}\mathbf{H}) - tr(\mathbf{H}^{-1}) &= \frac{1}{\sigma_e^2}(\mathbf{y}^\top \mathbf{H}^{-1}\mathbf{y} - \mathbf{y}^\top \mathbf{H}^{-1}\mathbf{H}^{-1}\mathbf{y}) \\
n\sigma_e^2 - tr(\mathbf{H}^{-1})\sigma_e^2 &= \mathbf{y}^\top \mathbf{H}^{-1}\mathbf{y} - \mathbf{y}^\top \mathbf{H}^{-1}\mathbf{H}^{-1}\mathbf{y} \\
n\sigma_e^2 &= \mathbf{y}^\top \mathbf{H}^{-1}\mathbf{y} + tr(\mathbf{H}^{-1})\left(\sigma_e^2 - \frac{\mathbf{y}^\top \mathbf{H}^{-1}\mathbf{H}^{-1}\mathbf{y}}{tr(\mathbf{H}^{-1})}\right)
\end{aligned} \tag{6}$$

Now consider the score $U_{\sigma_e^2} = \mathbf{0}$. Using the same quantity $\mathbf{V}^{-1} = \frac{\mathbf{H}^{-1}}{\sigma_e^2}$, we have:

$$\begin{aligned}
tr(\mathbf{V}^{-1}) &= \mathbf{y}^\top \mathbf{V}^{-1}\mathbf{V}^{-1}\mathbf{y} \\
tr(\mathbf{H}^{-1}) &= \frac{\mathbf{y}^\top \mathbf{H}^{-1}\mathbf{H}^{-1}\mathbf{y}}{\sigma_e^2}
\end{aligned} \tag{7}$$

Plugging (7) into (6) and substituting $H$ back with $V$, we have:

$$\begin{aligned}
n\sigma_e^2 &= \mathbf{y}^\top \mathbf{H}^{-1}\mathbf{y} \\
\Rightarrow \sigma_e^2 &= \frac{\mathbf{y}^\top \mathbf{H}^{-1}\mathbf{y}}{n} = \frac{\mathbf{y}^\top \mathbf{V}^{-1}\mathbf{y}\sigma_e^2}{n}.
\end{aligned} \tag{8}$$

Finally, to express $\sigma_e^2$ as a function of BLUP, we use the the Woodbury formula to get:

$$
\begin{aligned}
\mathbf{V}^{-\mathbf{1}}\mathbf{y} &= \left(\frac{1}{\sigma_e^2}\mathbf{I} - \frac{1}{\sigma_e^4}\mathbf{X}\left(\frac{1}{\sigma_g^2}\mathbf{I} + \mathbf{X}^\top\mathbf{X}\frac{1}{\sigma_e^2}\right)^{-1}\mathbf{X}^\top\right)\mathbf{y} \\
&= \frac{1}{\sigma_e^2}\mathbf{y} - \frac{1}{\sigma_e^2}\mathbf{X}\left(\frac{1}{\sigma_g^2}\mathbf{I} + \mathbf{X}^\top\mathbf{X}\frac{1}{\sigma_e^2}\right)^{-1}\mathbf{X}^\top\mathbf{y}\frac{1}{\sigma_e^2} \\
&= \frac{1}{\sigma_e^2}\mathbf{y} - \frac{1}{\sigma_e^2}\mathbf{X}\left(\frac{\sigma_e^2}{\sigma_g^2}\mathbf{I} + \mathbf{X}^\top\mathbf{X}\right)^{-1}\mathbf{X}^\top\mathbf{y} \\
&= \frac{1}{\sigma_e^2}(\mathbf{y} - \mathbf{X}\tilde{\boldsymbol{\beta}}) \qquad (9)
\end{aligned}
$$

where $\tilde{\beta}$ is the BLUP estimate: $\left(\frac{\sigma_e^2}{\sigma_g^2}\mathbf{I} + \mathbf{R}\right)^{-1}\mathbf{S}$. Plugging (9) into (8) gives us the expression for updating $\sigma_e^2$ using the BLUP:

$$\sigma_e^2 = \frac{\mathbf{y}^\top\mathbf{y} - \mathbf{X}\tilde{\boldsymbol{\beta}}}{n}.$$

153 ### 3.3 Updating equation for $\sigma_g^2$

Define $\mathbf{W} = \left(\mathbf{I} + \frac{\sigma_g^2}{\sigma_e^2}\mathbf{R}\right)$. Then we have $\mathbf{W} - \mathbf{I} = \frac{\mathbf{R}\sigma_g^2}{\sigma_e^2}$. The LHS of the score equation can be rewritten, using the Woodbury identity matrix, we have:

$$
\begin{aligned}
tr(\mathbf{X}^\top\mathbf{V}^{-\mathbf{1}}\mathbf{X}) &= tr\left(\mathbf{X}^\top\left(\frac{1}{\sigma_e^2}\left(\mathbf{I} - \frac{\sigma_g^2}{\sigma_e^2}\mathbf{X}\mathbf{W}^{-\mathbf{1}}\mathbf{X}^\top\right)\right)\mathbf{X}\right) \\
&= tr\left(\frac{\mathbf{R}}{\sigma_e^2} - \frac{\sigma_g^2}{\sigma_e^2}\mathbf{R}\mathbf{W}^{-1}\frac{\mathbf{R}}{\sigma_e^2}\right) \\
&= tr\left(\left(\frac{\mathbf{R}\sigma_g^2}{\sigma_e^2} - \frac{\mathbf{R}\sigma_g^2}{\sigma_e^2}\mathbf{W}^{-1}\frac{\mathbf{R}\sigma_g^2}{\sigma_e^2}\right)\frac{1}{\sigma_g^2}\right) \\
&= tr\left(\frac{1}{\sigma_g^2}(\mathbf{W} - \mathbf{I} - (\mathbf{W} - \mathbf{I})\mathbf{W}^{-1}(\mathbf{W} - \mathbf{I}))\right) \\
&= tr\left(\frac{1}{\sigma_g^2}(\mathbf{I} - \mathbf{W}^{-1})\right) \\
&= \frac{p - tr(\mathbf{W}^{-1})}{\sigma_g^2}. \qquad (10)
\end{aligned}
$$

The RHS of the score equation can be rewritten as:

$$\mathbf{y}^\top \mathbf{V}^{-1}\mathbf{X}\mathbf{X}^\top \mathbf{V}^{-1}\mathbf{y} = \frac{\mathbf{y}^\top \mathbf{V}^{-1}\mathbf{X}\sigma_g^2\sigma_g^2\mathbf{X}^\top \mathbf{V}^{-1}\mathbf{y}}{\sigma_g^4} = \frac{\tilde{\boldsymbol{\beta}}^\top \tilde{\boldsymbol{\beta}}}{\sigma_g^4}. \tag{11}$$

Equating (10) and (11) leads us to have two equivalent updating equations for $\sigma_g^2$:

$$\sigma_g^2 = \frac{\tilde{\boldsymbol{\beta}}^\top \tilde{\boldsymbol{\beta}} + \sigma_e^2 tr(\mathbf{W}^{-1})}{p} \quad \text{or} \quad \sigma_g^2 = \frac{\tilde{\boldsymbol{\beta}}^\top \tilde{\boldsymbol{\beta}}}{p - tr(\mathbf{W}^{-1})}.$$

In summary, we have shown that the score equations of $(\sigma_g^2, \sigma_e^2)$ derived from the likelihood in (1) are identical to the score equations based on (5), giving rise to our updating equations in the HEELS estimation procedure.

#### 4 HEELS estimation with unknown sample variance

When sample variance is not known, we can approximate $\mathbf{y}^\top\mathbf{y}/n$ in two ways. Suppose the effect sizes and standard errors are both provided in the summary statistics. For SNP $j$, consider the marginal model used in the association studies, $\mathbf{y} = \mathbf{X}_j\beta^j + \varepsilon^j$, where we assume $\mathbf{X}_j$ has been standardized to have variance 1, for all $j$. Superscripts are used to distinguish the models for different markers while subscripts indicate specific columns of the genotype matrix.

$$\begin{aligned}
\frac{\mathbf{y}^\top\mathbf{y}}{n} &= \frac{1}{n}(\mathbf{X}_j\beta^j + \varepsilon^j)^\top(\mathbf{X}^j\beta^j + \varepsilon^j) \\
&= \frac{1}{n}(\beta_j^2\mathbf{X}_j^\top\mathbf{X}_j + 2\beta^j\mathbf{X}_j^\top\varepsilon^j + [\varepsilon^j]^\top\varepsilon^j) \\
&= \beta_j^2 + \frac{[\varepsilon^j]^\top\varepsilon^j}{n} \\
&= \beta_j^2 + [\sigma_e^j]^2 \approx \frac{1}{p}\sum_{j=1}^{p}([\beta^j]^2 + n[SE^j]^2),
\end{aligned}$$

where we use the mean estimator in the last step. Alternatively, we can assume a unit-variance phenotype and use the standardized effect sizes to estimate $\hat{h}^2_{HEELS}$. We use this second approximation in practice, as it is more generally applicable, especially when only the Z-scores or $p$-values are provided in the summary statistics. Accordingly, we normalize the variance component estimates at each iteration such that $\sigma_g^2 + \sigma_e^2 = 1$.

#### 5 HEELS as an EM algorithm

In this section, we demonstrate that the HEELS procedure can be viewed as an EM algorithm. We proceed by first formulating the EM algorithm using the same model as the one we used for deriving HEELS. Then we connect it to the HEELS estimation procedure by showing its equivalence to the REML estimating equations. In essence, EM maximizes the likelihood of the complete or augmented data, which in our case is the phenotype vector and the unobserved random effects, $(\mathbf{y}, \beta)$. The incomplete data, on the other hand, is the observed phenotype vector, $\mathbf{y}$.

EM iterates between updating the random effects using its expected conditional mean, and updating the variance components using the maximizers of the log-likelihood based on the joint density. Given the joint distribution of the complete data specified in equation (4), the conditional distribution of the random effects is:

$$\beta|\mathbf{y} \sim MVN(\sigma_g^2\mathbf{X}^\top\mathbf{V}^{-\mathbf{1}}\mathbf{y}, \sigma_g^2\mathbf{I}_p - \sigma_g^4\mathbf{X}^\top\mathbf{V}^{-\mathbf{1}}\mathbf{X}).$$

Hence, the conditional expected value of the random effects is $\mathbb{E}(\beta|\mathbf{y}) = \sigma_g^2\mathbf{X}^\top\mathbf{V}^{-\mathbf{1}}\mathbf{y}$. The log-likelihood based on the complete data is,

$$l(\sigma_g^2, \sigma_e^2; \mathbf{y}, \boldsymbol{\beta}) \propto -\frac{1}{2}\left\{p \cdot log(\sigma_g^2) + n \cdot log(\sigma_e^2) + \frac{\boldsymbol{\beta}^\top\boldsymbol{\beta}}{\sigma_g^2} + \frac{(\mathbf{y}-\mathbf{X}\boldsymbol{\beta})^\top(\mathbf{y}-\mathbf{X}\boldsymbol{\beta})}{\sigma_e^2}\right\}, \tag{12}$$

and the maximizers of which can be easily derived as,

$$\sigma_g^2 = \frac{\boldsymbol{\beta}^\top\boldsymbol{\beta}}{p} \quad \text{and} \quad \sigma_e^2 = \frac{(\mathbf{y}-\mathbf{X}\beta)^\top(\mathbf{y}-\mathbf{X}\beta)}{n}. \tag{13}$$

The maximization step uses the conditional expectation $\mathbb{E}(\beta|\mathbf{y})$ in place of the true unknown random effects $\beta$ in (13). To summarize, the EM procedure for estimating the variance components alternates between the following two steps,

- At the E-step, we update the random effects using the incomplete data and the current value of the variance components from the M-step,

$$\boldsymbol{\beta}^{(t)} = \sigma_g^{2(t)}\mathbf{X}^\top\mathbf{V}^{-\mathbf{1}}\mathbf{y}. \tag{14}$$

- At the M-step, we update the variance components using the complete data and the conditional expectation of the random effects from the E-step,

$$\sigma_g^{2(t)} = \frac{\mathbb{E}(\boldsymbol{\beta}^\top \boldsymbol{\beta}|\mathbf{y})}{p}\bigg|_{\beta^{(t)}} = \frac{1}{p}\left(\sigma_g^{4(t-1)}\mathbf{y}^\top\mathbf{V}^{-\mathbf{1}}\mathbf{X}\mathbf{X}^\top\mathbf{V}^{-\mathbf{1}}\mathbf{y} + tr(\sigma_g^{2(t-1)}\mathbf{I}_p - \sigma_g^{4(t-1)}\mathbf{X}^\top\mathbf{V}^{-\mathbf{1}}\mathbf{X})\right) \tag{15}$$

$$\sigma_e^{2(t)} = \frac{\mathbb{E}((\mathbf{y}-\mathbf{X}\boldsymbol{\beta})^\top(\mathbf{y}-\mathbf{X}\boldsymbol{\beta})|\mathbf{y})}{n}\bigg|_{\beta^{(t)}} = \frac{1}{n}\left(\sigma_e^{4(t-1)}\mathbf{y}^\top\mathbf{V}^{-\mathbf{2}}\mathbf{y} + tr(\sigma_e^{2(t-1)}\mathbf{I}_n - \sigma_e^{4(t-1)}\mathbf{V}^{-1})\right) \tag{16}$$

where the second equality in both equation (15) and (16) follows from the conditional distributional form of $\beta|\mathbf{y}$ and the quadratic form theory.

To see the connections between the EM algorithm and our estimation procedure, note that the updating equations (15) and (16) are fixed-point iterations which lead to the same solutions as the estimating equations solved by HEELS.

$$\begin{aligned}
p\sigma_g^2 &= \sigma_g^4\mathbf{y}^\top\mathbf{V}^{-\mathbf{1}}\mathbf{X}\mathbf{X}^\top\mathbf{V}^{-\mathbf{1}}\mathbf{y} + tr(\sigma_g^2\mathbf{I}_p - \sigma_g^4\mathbf{X}^\top\mathbf{V}^{-\mathbf{1}}\mathbf{X}) \\
0 &= \mathbf{y}^\top\mathbf{V}^{-\mathbf{1}}\mathbf{X}\mathbf{X}^\top\mathbf{V}^{-\mathbf{1}}\mathbf{y} - tr(\mathbf{X}^\top\mathbf{V}^{-\mathbf{1}}\mathbf{X}) && \text{(same as equation (6))} \\
n\sigma_e^2 &= \sigma_e^4\mathbf{y}^\top\mathbf{V}^{-\mathbf{2}}\mathbf{y} + tr(\mathbf{I}_n - \sigma_e^2\mathbf{V}^{-1}) \\
0 &= \mathbf{y}^\top\mathbf{V}^{-\mathbf{2}}\mathbf{y} - tr(\mathbf{V}^{-1}). && \text{(same as equation (7))}
\end{aligned}$$

Lastly, the updating equation in (14) coincide with the updating equation for $\beta$ in HEELS. Therefore, the HEELS estimation procedure can be viewed as an EM algorithm under the same model. Notably, HEELS has several nice properties which are also features of the EM algorithm, such as guarantee of convergence under fairly unrestricted conditions[34,35]. Viewing the HEELS algorithm as an EM is a helpful step for extending our method to incorporate sparse or heterogeneous effect size variances. For example, we can modify the complete likelihood function to incorporate flexible weighting of SNPs; moreover, we can introduce latent variables to indicate the markers' null vs non-null effects and expand the E-step to update this latent variable together with $\boldsymbol{\beta}$.

#### 6 Relationship between GRE and HESS

Here we explain the similarities and differences between the GRE estimator and the HESS estimator of SNP-heritability. First of all, both the GRE estimator and the HESS estimator have a closed-form expression and are functions of the marginal association statistics and in-sample LD. While GRE is a genome-wide $h^2_{SNP}$ estimator, it assumes independent and additive chromosome-wide heritabilities. Hence, the form of the *regional* GRE estimator in fact coincides with that of the HESS estimator when LD is estimated in-sample,

$$\hat{h}^2_{GRE} = \sum_{c=1}^{22} \frac{n\hat{\beta}_c^\top \mathbf{R}_c^\dagger \hat{\beta}_c - p_c}{n - p_c} \quad \text{, where } c \text{ signifies chromosom;}$$

$$\hat{h}^2_{HESS_k} = \frac{n\hat{\beta}_k^\top \mathbf{R}_k^\dagger \hat{\beta}_k - p_k}{n - p_k} \quad \text{, where } k \text{ signifies region.}$$

$\mathbf{R}_c^\dagger$ and $\mathbf{R}_k^\dagger$ are the pseudo-inverse of the empirical chromosome-wide or regional LD matrices.

GRE and HESS can both be viewed as principal-component regression (PCR) based estimators. In other words, both of these two estimators are weighted summations of the squared projections of GWAS effect sizes onto the eigenvectors of the LD matrix. Denote the singular value decomposition of $\mathbf{X}$ by $\mathbf{X} = \mathbf{U}\Lambda\mathbf{V}^\top$. Then the covariance matrix can be written as $\mathbf{X}^\top\mathbf{X} = \mathbf{V}\Lambda^2\mathbf{V}^\top$. We can project $\mathbf{X}$ onto a lower-dimensional space using the eigenvectors of its covariance matrix:

$$\mathbf{y} = \mathbf{X}\beta_J + \varepsilon = \underbrace{\mathbf{X}\mathbf{V}}_{\mathbf{Z}}\underbrace{\mathbf{V}^\top\beta_J}_{\gamma} + \varepsilon = \mathbf{Z}\gamma + \varepsilon,$$

where we use $\beta_J$ to denote the true joint effect size, to be distinguished from the marginal effect size $\beta_M = \mathbf{R}\beta_J$. Assuming unit-variance phenotypes or standardized effect sizes, the PCR-estimator for heritability when the top $r$ principal components are used is,

$$\hat{h}^2_{PCR_r} = 1 - \hat{\sigma}_e^2 = 1 - \frac{(\mathbf{y} - \mathbf{Z_r}\hat{\gamma}_\mathbf{r})^\top(\mathbf{y} - \mathbf{Z_r}\hat{\gamma}_\mathbf{r})}{n - r},$$

where the projected genotype matrix and effect size are $\mathbf{Z}_r = \mathbf{X}\mathbf{V_r}$ and $\gamma_r = \mathbf{V}_r^\top\beta_J$. Notably,

$$
\begin{aligned}
\hat{\sigma}_e^2 &= \frac{(\mathbf{y}-\mathbf{Z_r}\hat{\gamma}_\mathbf{r})^\top(\mathbf{y}-\mathbf{Z_r}\hat{\gamma}_\mathbf{r})}{n-r} \\
&= \frac{\mathbf{y}^\top\mathbf{y}-2\mathbf{y}^\top\mathbf{Z_r}\gamma_r+\gamma_r^\top\mathbf{Z_r}^\top\mathbf{Z_r}\gamma_r}{n-r} \\
&= \frac{\mathbf{y}^\top\mathbf{y}-2\mathbf{y}^\top\mathbf{X}\mathbf{V_r}\mathbf{V_r}^\top\beta_J+\beta_J^\top\mathbf{V_r}\mathbf{V_r}^\top\mathbf{X}^\top\mathbf{X}\mathbf{V_r}\mathbf{V_r}^\top\beta_J}{n-r} \\
&= \frac{n-2\beta_J^\top\mathbf{R}\mathbf{V_r}\mathbf{V_r}^\top\beta_J+\beta_J^\top\mathbf{R}\beta_J}{n-r} && \text{(orthogonality of } \mathbf{V}_r\text{)} \\
&= \frac{n-\beta_J^\top\mathbf{R}\beta_J}{n-r} \\
&= \frac{n-\beta_M^\top\mathbf{R}^{-1}\mathbf{R}n\mathbf{R}^{-1}\beta_M}{n-r} && \text{(use } \beta_J=\Sigma^{-1}\beta_M\text{)} \\
&= \frac{n-n\beta_M^\top\mathbf{R}^{-1}\beta_M}{n-r} \\
1-\hat{\sigma}_e^2 &= \frac{n\beta_M^\top\mathbf{R}^{-1}\beta_M-r}{n-r} && (17)
\end{aligned}
$$

Therefore, $\hat{h}^2_{PCR_r}$ is precisely the GRE estimator when $r=\min(n,p)=rank(\mathbf{X})$ and is the HESS estimator with truncated SVD using the top $r$ eigenvectors, when $\Sigma^{-1}$ is approximated by $\mathbf{V}_r\Lambda_r^{-2}\mathbf{V_r}^\top$.

An important similarity between the GRE estimator and the HESS estimator is that they are both robust to different underlying genetic architectures. HESS is unbiased as it models the causal effects as fixed; GRE is consistent as it allows arbitrary and SNP-specific variance of causal effects. Moreover, both methods require $n >> p$. The HESS estimator is applied at the local level to independent loci, so it does not require ultra-large sample size. The GRE estimator requires biobank-scale data as it is applied to the whole genome. For both estimators, the requirement for large sample size arises from the asymptotics used in deriving the unbiased estimator. The variance of both estimators are similarly developed based on quadratic form theory.

The two estimators differ in several aspects. First, GRE is derived under the framework of random effects whereas HESS assumes the genetic effects are fixed. In other words, the estimands or the target quantity $h^2_{SNP}$ defined by these two methods are slightly different:

$$\text{Random effects model (adopted by GRE): } h^2_{SNP}=\mathbb{E}(\beta^\top\Sigma\beta)=\sum_{j=1}^{p}\sigma_j^2;$$

$$\text{Fixed effects model (adopted by HESS): } h^2_{SNP}=\beta^\top\Sigma\beta.$$

where $\Sigma$ is the population LD and $\sigma_j^2$ is the marker-specific effect size variance for SNP $j$, assumed to be any arbitrary positive value. GRE can be viewed as a generalized method-of-moment estimator, as it was derived by equating the population and the sample moment $\beta_M^\top \mathbf{R}^{-1}\beta_M$ (see equation (4) in the Supplementary notes of Hou et al.[20] and the expression right above it). Note that the moment equating step assumes conditioning both of the effect size and genotype, the estimator may be considered as a fixed-effect heritability estimator.

Second, GRE uses in-sample LD information and thus does not require regularization of the LD matrix, whereas HESS is designed to account for the LD structure using a reference panel and thus regularizes out-of-sample LD via truncated SVD. When LD is estimated in-sample and no LD regularization is applied, *i.e.* $r = rank(\mathbf{X})$, the two estimators coincide completely. Indeed, we observed in simulations that the estimates from HESS approach the estimates from GRE as LD regularization lessens (**Supplementary Figure** 5).

### 7 The asymptotic variance of HEELS

We start by first deriving the information matrix for $\ell(\mathbf{y};\sigma_g^2,\sigma_e^2)$. Given that $U_{\sigma_g^2}(\mathbf{y}) = -\frac{1}{2}tr(\mathbf{X}^\top\mathbf{V}^{-\mathbf{1}}\mathbf{X}) + \frac{1}{2}(\mathbf{X}^\top\mathbf{V}^{-\mathbf{1}}\mathbf{y})^\top(\mathbf{X}^\top\mathbf{V}^{-\mathbf{1}}\mathbf{y})$, we have,

$$
\begin{aligned}
-\mathbb{E}\left(\frac{\partial U_{\sigma_g^2}(\mathbf{y})}{\partial \sigma_g^2}\right) &= -\frac{1}{2}tr(\mathbf{V}^{-\mathbf{1}}\mathbf{X}\mathbf{X}^\top\mathbf{V}^{-\mathbf{1}}\mathbf{X}\mathbf{X}^\top) + \mathbb{E}(\mathbf{y}^\top\mathbf{V}^{-\mathbf{1}}\mathbf{X}\mathbf{X}^\top\mathbf{V}^{-\mathbf{1}}\mathbf{X}\mathbf{X}^\top\mathbf{V}^{-\mathbf{1}}\mathbf{y}) \\
&= -\frac{1}{2}tr(\mathbf{V}^{-\mathbf{1}}\mathbf{X}\mathbf{X}^\top\mathbf{V}^{-\mathbf{1}}\mathbf{X}\mathbf{X}^\top) + tr(\mathbf{V}^{-\mathbf{1}}\mathbf{X}\mathbf{X}^\top\mathbf{V}^{-\mathbf{1}}\mathbf{X}\mathbf{X}^\top\mathbf{V}^{-\mathbf{1}}\mathbf{V}) \\
&= \frac{1}{2}tr(\mathbf{V}^{-\mathbf{1}}\mathbf{X}\mathbf{X}^\top\mathbf{V}^{-\mathbf{1}}\mathbf{X}\mathbf{X}^\top).
\end{aligned}
$$

Given that $U_{\sigma_e^2}(\mathbf{y}) = -\frac{1}{2}tr(\mathbf{V}^{-\mathbf{1}}) + \frac{1}{2}\mathbf{y}^\top\mathbf{V}^{-\mathbf{1}}\mathbf{V}^{-\mathbf{1}}\mathbf{y}$, we have,

$$
\begin{aligned}
-\mathbb{E}\left(\frac{\partial U_{\sigma_e^2}(\mathbf{y})}{\sigma_e^2}\right) &= -\frac{1}{2}tr(\mathbf{V}^{-2}) + \mathbb{E}(\mathbf{y}^\top\mathbf{V}^{-2}\mathbf{V}^{-\mathbf{1}}\mathbf{y}) \\
&= -\frac{1}{2}tr(\mathbf{V}^{-2}) + tr(\mathbf{V}^{-2}\mathbf{V}^{-\mathbf{1}}\mathbf{V}) \\
&= \frac{1}{2}tr(\mathbf{V}^{-2}).
\end{aligned}
$$

The cross-term of the information matrix can be derived as:

$$
\begin{aligned}
-\mathbb{E}\left(\frac{\partial U_{\sigma_g^2}(\mathbf{y})}{\partial \sigma_e^2}\right) &= -\frac{1}{2}tr(\mathbf{V}^{-1}\mathbf{X}\mathbf{X}^\top\mathbf{V}^{-1}) + \mathbb{E}(\mathbf{y}^\top\mathbf{V}^{-1}\mathbf{X}\mathbf{X}^\top\mathbf{V}^{-1}\mathbf{V}^{-1}\mathbf{y}) \\
&= -\frac{1}{2}tr(\mathbf{V}^{-1}\mathbf{X}\mathbf{X}^\top\mathbf{V}^{-1}) + tr(\mathbf{V}^{-1}\mathbf{X}\mathbf{X}^\top\mathbf{V}^{-1}\mathbf{V}^{-1}\mathbf{V}) \\
&= \frac{1}{2}tr(\mathbf{V}^{-1}\mathbf{X}\mathbf{X}^\top\mathbf{V}^{-1}).
\end{aligned}
$$

Therefore, for true values of the variance components $\sigma_e^2, \sigma_g^2$ and positive definite $\mathbf{V}$, the information matrix is:

$$
I(\sigma_e^2, \sigma_g^2; \mathbf{X}, \mathbf{y}) = \frac{1}{2}\begin{bmatrix} tr(\mathbf{V}^{-2}) & tr(\mathbf{V}^{-1}\mathbf{X}\mathbf{X}^\top\mathbf{V}^{-1}) \\ tr(\mathbf{V}^{-1}\mathbf{X}\mathbf{X}^\top\mathbf{V}^{-1}) & tr(\mathbf{V}^{-1}\mathbf{X}\mathbf{X}^\top\mathbf{V}^{-1}\mathbf{X}\mathbf{X}^\top) \end{bmatrix}.
$$

Using properties of trace and the identity of $\mathbf{W} := \frac{\sigma_e^2}{\sigma_g^2}\mathbf{I} + \mathbf{R}$[33], we can rewrite the information matrix using only the summary statistics and in-sample LD as the following:

$$
I(\sigma_e^2, \sigma_g^2; \mathbf{S}, \mathbf{R}) = \frac{1}{2}\begin{bmatrix} \frac{n-p}{\sigma_e^4} + \frac{1}{\sigma_g^4}tr(\mathbf{W}^{-2}) & \frac{1}{\sigma_g^4}tr(\mathbf{W}^{-1}) - \frac{\sigma_e^2}{\sigma_g^6}tr(\mathbf{W}^{-2}) \\ \frac{1}{\sigma_g^4}tr(\mathbf{W}^{-1}) - \frac{\sigma_e^2}{\sigma_g^6}tr(\mathbf{W}^{-2}) & \frac{p}{\sigma_g^4} - \frac{2\sigma_e^2}{\sigma_g^6}tr(\mathbf{W}^{-1}) + \frac{\sigma_e^4}{\sigma_g^8}tr(\mathbf{W}^{-2}) \end{bmatrix} \quad (18)
$$

### 8 Approximating the asymptotic variance of $h^2_{HEELS}$ with a low-dimensional representation of the LD

In this section, we provide details on the asymptotic variance of $h^2_{HEELS}$ when the LD matrix has a low dimensional representation. We only describe the scenario where the Banded + LR structure is employed, as the other strategies are special cases of this general setting. Let $\mathbf{W} := \frac{\sigma_e^2}{\sigma_g^2}\mathbf{I} + \mathbf{R}$ be the working matrix, as defined before. Suppose we have obtained the approximation form of the LD matrix, denoted by $\mathbf{R} \approx \mathbf{R}_b + \mathbf{U}_r\Lambda_r\mathbf{U}^\top$, we then have an approximation form of the working matrix $\mathbf{W} \approx \mathbf{W}_b + \mathbf{U}_r\Lambda_r\mathbf{U}^\top$ with $\mathbf{W}_b = \mathbf{R}_b + \frac{\sigma_e^2}{\sigma_g^2}I$. For any given matrices $\mathbf{Y} \in \mathbb{R}^{p\times p}, \mathbf{Z} \in \mathbb{R}^{p\times r}, \Lambda \in \mathbb{R}^{r\times r}$, we define a mapping,

$$
f : \mathbb{R}^{p\times p} \times \mathbb{R}^{p\times r} \times \mathbb{R}^{r\times r} \longrightarrow \mathbb{R}^{p\times p}
$$

$$
f(\mathbf{Y}, \mathbf{Z}, \Lambda) = \mathbf{Y} - \mathbf{Y}\mathbf{Z}(\Lambda^{-1} + \mathbf{Z}^\top\mathbf{Y}\mathbf{Z})^{-1}\mathbf{Z}^\top\mathbf{Y}^\top.
$$

This mapping corresponds to the Woodbury matrix inverse formula, *i.e.* $\mathbf{W}^{-1} = f(\mathbf{W}_b, \mathbf{U}_r, \Lambda_r)$ if $\mathbf{W} = \mathbf{W}_b + \mathbf{U}_r\Lambda_r\mathbf{U}^\top$. We can apply this mapping again to express the squared term, $\mathbf{W}^{-2}$,

$$\begin{aligned}
\mathbf{W}^{-2} &\approx (\mathbf{W}_b^\top\mathbf{W}_b + 2\mathbf{W}_b^\top\mathbf{U}_r\Lambda_r\mathbf{U}_r^\top + \mathbf{U}_r\Lambda_r^2\mathbf{U}_r^\top)^{-1} \\
&= (\mathbf{V} + \mathbf{U}_r\Lambda_r^2\mathbf{U}_r^\top)^{-1} \qquad (\text{Let } \mathbf{V} = \mathbf{W}_b^\top\mathbf{W}_b + 2\mathbf{W}_b^\top\mathbf{U}_r\Lambda_r\mathbf{U}_r^\top) \\
&\approx \mathbf{V}^{-1} - \mathbf{V}^{-1}\mathbf{U}_r(\Lambda_r^{-2} + \mathbf{U}_r^\top\mathbf{V}^{-1}\mathbf{U}_r)^{-1}\mathbf{U}_r^\top\mathbf{V}^{-1} \\
&= f(\mathbf{V}^{-1}, \mathbf{U}_r, \Lambda_r^2).
\end{aligned}$$

Furthermore, we note that $\mathbf{V}^{-1} = (\mathbf{W}_b + 2\mathbf{U}_r\Lambda_r\mathbf{U}_r^\top)^{-1}\mathbf{W_b}^{-1} = f(\mathbf{W}_b, \mathbf{U}_r, 2\Lambda_r)\mathbf{W_b}^{-1}$. Hence, both $\mathbf{W}^{-1}$ and $\mathbf{W}^{-2}$ can be expressed as functions of the approximating elements of the LD matrix:

$$\mathbf{W}^{-1} = f(\mathbf{W}_b, \mathbf{U}_r, \Lambda_r) \tag{19}$$

$$\mathbf{W}^{-2} = f(f(\mathbf{W}_b, \mathbf{U}_r, 2\Lambda_r)\mathbf{W}_b^{-1}, \mathbf{U}_r, \Lambda_r^2). \tag{20}$$

Therefore, in cases where the LD matrix has been approximated by the sum of a banded matrix and a low-rank matrix, we can still estimate the variance of the HEELS estimator without incurring additional computational costs.

### 9 Theoretical comparison of statistical efficiency between HEELS and LDSC

We demonstrate how our estimator is equivalent to the most efficient estimator under the Generalized Method of Moments (GMM) framework. Given the same modeling of the phenotypes as is defined in the main text, but without the distributional assumption, the estimating equations for the method of moments estimators take the form of,

$$\mathbf{y}^\top\mathbf{A}_1\mathbf{y} = tr(\mathbf{A}_1\mathbf{X}\mathbf{X}^\top)\sigma_g^2 + tr(\mathbf{A}_1)\sigma_e^2 \tag{21}$$

$$\mathbf{y}^\top\mathbf{A}_2\mathbf{y} = tr(\mathbf{A}_2\mathbf{X}\mathbf{X}^\top)\sigma_g^2 + tr(\mathbf{A}_2)\sigma_e^2, \tag{22}$$

where $\mathbf{A_1}, \mathbf{A_2}$ are two symmetric non-negative definite matrices of dimension $n \times n$, used as weights. From the theory of method of moments, the choice of $\mathbf{A}$ will not affect the unbiasedness of $\sigma_g^2, \sigma_e^2$, but can affect the statistical efficiency of the estimators. A common criterion for selecting the optimal weight matrices is to minimize the expected squared error (*i.e.* the difference between population and sample moments), which is called the "Minimal Norm Quadratic Unbiased Estimation (MINQUE)" criterion[19]. For $l = 1,2$, we solve the following objective function,

$$\min E\left[\left(\mathbf{y}^\top \mathbf{A_l y} - tr(\mathbf{A}_l \mathbf{X}\mathbf{X}^\top)\sigma_g^2 - tr(\mathbf{A}_l)\sigma_e^2\right)^2\right], \tag{23}$$

which leads to the Best Quadratic Unbiased Estimator (BQUE). The optimal weights that minimize the objective in (23) are:

$$\mathbf{A}_1^* = \mathbf{V}^{-1}\mathbf{X}\mathbf{X}^\top\mathbf{V}^{-\mathbf{1}}; \quad \mathbf{A}_2^* = \mathbf{V}^{-\mathbf{1}}\mathbf{V}^{-\mathbf{1}}.$$

Since both $\mathbf{A}_1^*$ and $\mathbf{A}_2^*$ involve unknown population parameters through $\mathbf{V}$, the estimators of $\sigma_g^2, \sigma_e^2$ can only be obtained via an iterative procedure[19]. We next demonstrate how our HEELS estimator minimizes the expected squared error and thus meets the MINQUE criterion. It suffices to show that our score equations (2) and (3) coincide with the moment matching equations (21) and (22). With the optimal weights, $\mathbf{A}_1^*, \mathbf{A}_2^*$, equation (21) can be equated with equation (2) as:

$$\begin{aligned}
\mathbf{y}^\top(\mathbf{V}^{-\mathbf{1}}\mathbf{X}\mathbf{X}^\top\mathbf{V}^{-\mathbf{1}})\mathbf{y} &= tr(\mathbf{V}^{-\mathbf{1}}\mathbf{X}\mathbf{X}^\top\mathbf{V}^{-\mathbf{1}}\mathbf{X}\mathbf{X}^\top)\sigma_g^2 + tr(\mathbf{V}^{-\mathbf{1}}\mathbf{X}\mathbf{X}^\top\mathbf{V}^{-\mathbf{1}})\sigma_e^2 \\
(\mathbf{X}^\top\mathbf{V}^{-\mathbf{1}}\mathbf{y})^\top(\mathbf{X}^\top\mathbf{V}^{-\mathbf{1}}\mathbf{y}) &= tr(\mathbf{V}^{-\mathbf{1}}\mathbf{X}\mathbf{X}^\top\mathbf{V}^{-\mathbf{1}}(\mathbf{X}\mathbf{X}^\top\sigma_g^2\mathbf{I}_n + \sigma_e^2\mathbf{I}_n)) \\
&= tr(\mathbf{V}^{-\mathbf{1}}\mathbf{X}\mathbf{X}^\top\mathbf{V}^{-\mathbf{1}}\mathbf{V}) \\
&= tr(\mathbf{X}^\top\mathbf{V}^{-\mathbf{1}}\mathbf{X}).
\end{aligned}$$

Analogously, equation (22) can be equated with equation (3) as:

$$\begin{aligned}
\mathbf{y}^\top(\mathbf{V}^{-\mathbf{1}}\mathbf{V}^{-\mathbf{1}})\mathbf{y} &= tr(\mathbf{V}^{-\mathbf{1}}\mathbf{V}^{-\mathbf{1}}\mathbf{X}\mathbf{X}^\top)\sigma_g^2 + tr(\mathbf{V}^{-\mathbf{1}}\mathbf{V}^{-\mathbf{1}})\sigma_e^2 \\
(\mathbf{V}^{-\mathbf{1}}\mathbf{y})^\top(\mathbf{V}^{-\mathbf{1}}\mathbf{y}) &= tr(\mathbf{V}^{-\mathbf{1}}\mathbf{V}^{-\mathbf{1}}(\mathbf{K}\boldsymbol{\sigma}_{\mathbf{g}}^{\mathbf{2}}\mathbf{I}_n + \sigma_e^2\mathbf{I}_n)) \\
&= tr(\mathbf{V}^{-\mathbf{1}}).
\end{aligned}$$

Zhou (2017) showed that the heritability estimator of LDSC (without or with known population structure) can be viewed as a generalized method of moments estimator under the GMM framework, but with sub-optimal weights[19]. In other word, the statistical efficiency of LDSC is lower than that of BQUE. As we have established the equivalence of efficiency between our HEELS estimator and the BQUE above, we conclude that HEELS is statistically more efficient than LDSC, as is confirmed by the simulation results.

#### 10 The PSD assumption in low-dimensional representation of LD

In two of our proposed "Banded + LR" LD approximation strategies, "Seq_Band_LR" and "PSD_Band_LR", we impose the PSD assumption, which helps to both 1) reduce the computational burden of solving the LD decomposition, and 2) improve the efficiency of our heritability estimation algorithm.

First, it can reduce the computational burden of searching for the low-dimensional representation. Consider the following two minimization problems,

$$\tilde{\mathbf{R}}^b, \tilde{\mathbf{R}}^r = \arg \min_{\mathbf{R}^b, \mathbf{R}^r \in \mathbb{R}^{p \times p}} ||\mathbf{R} - \mathbf{R}^b - \mathbf{R}^r||_F^2 \tag{24}$$

$$\tilde{\mathbf{L}}^b, \tilde{\mathbf{U}}^r = \arg \min_{\mathbf{L}^b, \mathbf{U}^r \in L_p(\mathbb{R})} ||\mathbf{R} - \mathbf{L}^{b\top}\mathbf{L}^b - \mathbf{U}^{r\top}\mathbf{U}^r||_F^2 \tag{25}$$

where $L_p(\mathbb{R})$ denotes the set of $p \times p$ lower triangular matrices with real entries. The objective of both of these optimization procedures is to minimize the squared Frobenius norm of the error matrix. However, optimizing over the Cholesky factors of the banded and the low-rank components of the representation in equation (25) leads to fewer parameters than optimizing all elements of the banded and low-rank matrices in equation (24). The reduction of computational burden is substantial when $p$ is large.

Another benefit of using the PSD assumption is that the Cholesky factors, $\tilde{\mathbf{L}}^b, \tilde{\mathbf{U}}^r$, can be directly used to compute the inverse of the LD matrix in the HEELS estimation procedure, without incurring additional cost. To be specific, the major computational burden of the HEELS estimating procedure lies in the evaluation of $\mathbf{W}^{(\mathbf{t})-\mathbf{1}}$ and $tr(\mathbf{W}^{(\mathbf{t})-\mathbf{1}})$. In the absence of LD approximation, we compute the Cholesky factors of the working matrix $\mathbf{W}^{(t)}$ once per iteration, and these factors are used to calculate $\mathbf{W}^{(\mathbf{t})-\mathbf{1}}\mathbf{S}$ and $tr(\mathbf{W}^{(\mathbf{t})-\mathbf{1}})$ respectively. The PSD guarantee of the solutions lessens the computational cost of these two steps significantly by circumventing the need to compute the Cholesky factors of the working matrix $\mathbf{W}^{(t)}$ at every iteration.

Suppose the LD matrix has been approximated as $\mathbf{R} \approx \mathbf{R}_b + \mathbf{U}_r \Lambda_r \mathbf{U}_r^\top$. We apply the Sherman-Morrison-Woodbury formula to circumvent the inversion of $\mathbf{W}^{(t)}$ as the following,

$$\mathbf{W}^{(t)-1}\mathbf{S} = \mathbf{W}^{b(t)-1}\mathbf{S} - \mathbf{W}^{b(t)-1}\mathbf{U}_r \left( \Lambda_r^{-1} + \mathbf{U}_r \mathbf{W}^{b(t)-1} \mathbf{U}_r \right)^{-1} \mathbf{U}_r^\top \mathbf{W}^{b(t)-1}\mathbf{S}, \tag{26}$$

where $\mathbf{W}^{b(t)} = \frac{\sigma_e^{2(t)}}{\sigma_g^{2(t)}}\mathbf{I} + \mathbf{R}^b$. We can easily compute the first term in equation (26) by taking advantage of its banded structure. We calculate the second term of equation (26) from right to left, keeping the order of matrix-vectir multiplication to $\mathscr{O}(pr^2)$ instead of $\mathscr{O}(p^3)$. Since $\left(\Lambda_r^{-1} + \mathbf{U}_r \mathbf{W}^{b(t)-1} \mathbf{U}_r\right)$ is low-rank, its inversion only costs $\mathscr{O}(r^3)$. Overall, we bring down the cost of computing $\mathbf{W}^{(t)-1}\mathbf{S}$ from $\mathscr{O}(p^3)$ to $\mathscr{O}(pr^3 + bp^2)$.

To compute $tr(\mathbf{W}^{(t)-1})$, we take the trace on both sides of the Woodbury identity for $\mathbf{W}^{(t)}$,

$$\begin{aligned} tr(\mathbf{W}^{(t)-1}) &= tr(\mathbf{W}^{b(t)-1}) - tr\left(\left(\mathbf{W}^{b(t)-1}\mathbf{U}_r v_r\right)^\top \left(\mathbf{W}^{b(t)-1}\mathbf{U}_r v_r\right)\right) \\ &= tr(\mathbf{W}^{b(t)-1}) - \left|\left|\mathbf{W}^{b(t)-1}\mathbf{U}_r v_r\right|\right|_F^2, \end{aligned} \tag{27}$$

where $v_r = \left(\Lambda_r^{-1} + \mathbf{U_r}^\top \mathbf{W}^{b(t)-1}\mathbf{U_r}\right)^{-\frac{1}{2}}$ and we re-write the trace-inverse term using matrix form. Since the quantity $\mathbf{W}^{b(t)-1}$ has been calculated in the former step (26), both of the two terms on the right-hand side can be easily obtained. Hence the cost of computing $tr(\mathbf{W}^{(t)-1})$ is reduced from $\mathscr{O}(p^3)$ to $\mathscr{O}(bp^2)$. We also avoid storing the full $p^2$ matrix from iteration to iteration, so that the memory cost of our algorithm is kept at $\mathscr{O}(p \min\{b, r\})$.

### 11 Algorithms for hyperparameter tuning

An important aspect of our low-dimensional LD representation algorithm is the tuning of the hyperparameters. While heuristics or prior knowledge about the structure of the LD can be used to determine the optimal values of $(b, r)$, we used a more principled way to evaluate the performance of a representation. We propose using a data-adaptive procedure to identify the best representation of the LD matrix, using synthetic phenotypic data and cross-validation (**Algorithm** 1). We also propose an incremental SVD algorithm to tune the hyperparameters more efficiently, where the number of low-rank factors increases step by step and the performance of the HEELS estimator is evaluated dynamically (**Algorithm** 2). The incremental SVD algorithm is applicable to "Seq_Band_LR" and "PSD_Band_LR" strategies.

To speed up the low-rank decomposition, we replaced the *exact* solutions based on direct eigen-decomposition with the *approximate* solutions, and compared two approaches – one is the optimization approach ("optim"), where we solve $\arg\min_{\mathbf{U}^r \in L_p(\mathbb{R})} ||\mathbf{R}_{resid} - \mathbf{U}^{r\top}\mathbf{U}^r||_F^2$, and $\mathbf{R}_{resid}$ is the residual off-banded component to be decomposed or approximated and $L_p(\mathbb{R})$ is the collection of $p \times p$ lower triangular matrices; the other approach is based on random sketching ("random"), where we decompose $\mathbf{R}_{resid}$ via SVD[36]. The simulation results indicate that using the optimization approach is less prone to bias and is more robust to variation in the hyperparameter values (**Supplementary Figure** 10). While we observe

**Algorithm 1:** Pseudocode for selecting hyperparamter values of $b, r$

**Data:** $\mathbf{X} \in \mathbb{R}^{n \times p}$

**Input** : Strategy (one of the strategies in Table 2),
$h^2$ (arbitrary value between 0 and 1, used for generating synthetic phenotypes),
$m$ (number of replicates for cross-validation),
$\text{tol}_b, \text{tol}_r$ (stopping criteria, $tol_b < tol_r$),
$s$ (step size, default value is 50),
$\lambda$ (minimum sparsity, default value is 0.1)

**Initialize** : $r = 0, b = 0, e_{CV} = 1$

**while** $e_{CV} > tol_b \ \& \ b < \lambda p$ **do**
  $b \longleftarrow b + s$;
  **while** $e_{CV} > tol_r \ \& \ r < \lambda p$ **do**
    $r \longleftarrow r + s$;
    $(\hat{\mathbf{R}}_b, \hat{\mathbf{U}}_r, \hat{\mathbf{\Lambda}}_r) \longleftarrow$ **HEELS_sparse_LD**($\mathbf{R}, b, r$, Strategy)
    **for** $i = 1, 2, ..., m$ **do**
      $\beta_i \sim N(0, h^2/p)$;   $y_i = \mathbf{X}\beta_i$;
      $\hat{h}_i^2 \longleftarrow$ **HEELS_estimate_**$h^2$($\mathbf{X}^\top y_i, \hat{\mathbf{R}}_b, \hat{\mathbf{U}}_r, \hat{\mathbf{\Lambda}}_r$)
    **end**
    $e_{CV} = \left|\frac{1}{m}\sum_{i=1}^{m} \hat{h}_i^2 - h^2\right|$; ;   /* Cross-validation error */
  **end**
**end**
$b^*, r^* \longleftarrow b, r$;
$\mathbf{R}_b^*, \mathbf{U}_r^*, \mathbf{\Lambda}_r^* \longleftarrow \hat{\mathbf{R}}_b, \hat{\mathbf{U}}_r, \hat{\mathbf{\Lambda}}_r$

**Output** : $b^*, r^*$ (optimal hyperparameter values),
$\mathbf{R}_b^*, \mathbf{U}_r^*, \Lambda_r^*$ (optimal representation)

larger bias when the randomized SVD approach is used to approximate the low-rank component, it can be more advantageous in applications with larger problem size.

**Algorithm 2:** Incremental SVD algorithm for selecting optimal $r$

**Data:** $\mathbf{R}^* = \mathbf{R} - \hat{\mathbf{R}}_b$ (residual after the banded component has been estimated)
**Input :** $tol$ (stopping criteria), $s$ (increment size), $\lambda$ (minimum sparsity)
**Initialize :** $r = 0, i = 0, \mathbf{R}_r^0 = \mathbf{0}$

```
while e > tol & r < λp do
    i ⟵ i + 1;
    r ⟵ r + s ;
    (Û_r^i.Λ̂_r^i) ⟵ argmin_{Λ_r,U_r} ||R_* − R_r^{i−1} − U_r Λ_r U_r^⊤||_F^2;
    R_r^i ⟵ R_r^{i−1} + Û_r^i Λ̂_r^i Û_r^{i⊤};
    e ⟵ ||R^* − R_r^{i−1}||_F / ||R^*||_F ;    /* Approximation error of the current low-rank
        representation */
end
r^* ⟵ r; U_r^* ⟵ ∑_i U_r^i; Λ_r^* ⟵ ∑_i Λ̂_r^i
```

where line 4 is $(\hat{\mathbf{U}}_r^i.\hat{\Lambda}_r^i) \longleftarrow \arg\min_{\Lambda_r,U_r} ||\mathbf{R}_* - \mathbf{R}_r^{i-1} - \mathbf{U}_r\Lambda_r\mathbf{U}_r^\top||_F^2$; line 5 is $\mathbf{R}_r^i \longleftarrow \mathbf{R}_r^{i-1} + \hat{\mathbf{U}}_r^i\hat{\Lambda}_r^i\hat{\mathbf{U}}_r^{i\top}$; line 6 is $e \longleftarrow \frac{||\mathbf{R}^*-\mathbf{R}_r^{i-1}||_F}{||\mathbf{R}^*||_F}$; line 8 is $r^* \longleftarrow r$; $\mathbf{U}_r^* \longleftarrow \sum_i \mathbf{U}_r^i$; $\Lambda_r^* \longleftarrow \sum_i \hat{\Lambda}_r^i$

**Output :** $r^*$ (optimal hyperparameter value), $\mathbf{U}_r^*$, $\Lambda_r^*$ (optimal low-rank representation)
